# Exploring the Correlation Between UVB Sensitivity and SLE Activity: Insights into UVB-Driven Pathogenesis in Lupus Erythematosus

**DOI:** 10.1101/2024.09.28.615513

**Authors:** Jiayu He, Yuanning Guo, Jiamin Chen, Jinhua Xu, Xiaohua Zhu

## Abstract

Lupus erythematosus (LE) comprises various autoimmune inflammatory diseases, with significant overlap between cutaneous LE (CLE) and systemic LE (SLE). A key feature of both CLE and SLE is UV photosensitivity, particularly in UV-exposure-related skin inflammation. Despite this, reliable and objective UVB photosensitivity indicators closely correlating with LE activity have yet to be identified, and the underlying cellular and molecular mechanisms linking UVB sensitivity with LE onset and progression remain unclear.

We discovered that ultraviolet B minimal erythema dose (UVB-MED), a quantitative photosensitivity measure, is a significant and independent risk factor for SLE activity, demonstrating a negative correlation with SLEDAI (r = -0.58, *P* < 0.0001). Comprehensive transcriptomic analyses of large-scale CLE and SLE samples revealed more pronounced and extensive UVB-response gene dysregulation in skin tissues compared to blood. Additionally, 14 lupus activity-correlated, UVB-response genes (UVBACGs) were identified, including eight interferon-stimulated genes (IRF7, ISG20, ISG15, IFI44, IFITM1, MX1, LY6E, OASL) and others (JUN, PTTG1, HLA-F, CAV1, HOPX, RPL3), with dysregulation evident in both skin and blood cells, particularly immunocytes. Conventional LE therapies were found to be associated with these genes and may potentially regulate them, thereby contributing to therapeutic effects. These findings highlight the role of UVB in triggering autoimmune inflammation in the skin, which may subsequently spread to systemic inflammation via immune cells and factors. UVBACGs play a critical role in this process and may serve as targets for precise LE therapies, providing insight into the link between UVB photosensitivity and LE progression.

## Introduction

Lupus erythematosus (LE) refers a broad range of autoimmune inflammatory diseases most commonly appearing in the form of skin lesions. Systemic lupus erythematosus (SLE) is the most prevalent and serious LE subtype, which can affect any organ and usually cause life-threatening systemic complications [1, 2]. Although cutaneous lupus erythematosus (CLE) is mainly confined to skin lesions, over 20% of patients progress to SLE within three years and have increasing risks of further disease progression [3-5]. Within this period, CLE patients develop mild systemic disease, without meeting full SLE criteria [6]. In fact, approximately 80% SLE patients can suffer from skin lesions (25% are CLE specific lesions) at any stage of the disease [2, 5, 7, 8]. SLE and CLE can also coexist from 5% to 90% depend on different CLE subtypes, making it challenging to distinguish the two independently [4, 7, 8]. Noteworthily, skin autoimmune inflammation serves as a critical breakthrough that links CLE and SLE together. Nevertheless, there has been limited research focused on thoroughly examining the inflammatory similarities and differences between CLE and SLE, as well as investigating the mechanisms underlying their potential interconversion.

Ultraviolet light (UV) of skin exposure from sunlight is one of the crucial factors that can trigger and aggravate both SLE and CLE [9-11]. Notably, sun exposure can trigger systemic symptoms from CLE [12]. UV radiation, especially of ultraviolet light B (UVB), can exacerbate not only skin lesions but also LE systemic manifestations (e.g., fatigue, arthritis, nephritis), clinically and experimentally [9, 10, 13, 14]. Nearly 70–80% of LE patients are suffering from UV photosensitivity and around 50% of patients experience years of polymorphic light eruption before the onset of LE [11-15]. UV sensitivity and UV-radiation-associated skin autoimmune inflammatory pathogenesis are common features of CLE and SLE [10].

Despite the fact that photosensitivity was already reported linked to LE in 1851 and later included in the 11th ACR criteria [16] for the classification of SLE, it is still only vaguely defined as skin rashes resulting from unusual reaction to sunlight by patients’ history or physicians’ observation [2, 12, 16, 17]. Thus, later updated criteria SLICC [18] and EULAR [19] gradually removed the items related to photosensitivity. Phototesting, an objective method reproducing specific skin erythema when exposing to standardized artificial spectrums of UV radiation, has been standardized over 30 years for research and clinical evaluation. Whereas, the phototesting for photosensitivity has not been widely applied to clinical practice [10, 12, 20, 21]. This may be partly due to the fact that no studies have yet explored an objective and reliable UVB photosensitivity indicator that has a close correlation with LE activity.

Also, in order to push UV photosensitivity into diagnostic and therapeutic application, there is an urgent need for more consistent, comprehensive, and systematic studies to demonstrate how UVB-triggered autoimmunity promotes the onset and progression of LE, and how it enhances LE activity.

Therefore, the present study applied UVB minimal erythema dose (UVB-MED) as a quantitative and objective indicator for UVB sensitivity to detect its exact association with SLE activity. Then, by analyzing large cohorts of transcriptomic datasets from clinical samples of CLE and SLE tissues/cells (a total of 5,918 samples), we validated hundreds of UV-responsive candidate genes collected through systematic literature review, organization, and documentation. By comprehensively screening, identifying, and analyzing lupus activity-correlated, UVB-response genes (UVBACGs), we aim to interpret the relationship between UVB photosensitivity and LE initiation/aggression from the perspective of cellular and molecular mechanisms.

## Materials and Methods

### Clinical information collection, and criteria of disease activity and UVB-MED

We collected clinical data from consecutive SLE inpatients with UVB-MED phototesting records, admitted between July 2011 and December 2017, at the Department of Dermatology, Huashan Hospital. The inclusion and exclusion criteria for the patients were: 1) compliance with the American College of Rheumatology (ACR) 1997 [16] and the SLE International Clinical Cooperation group (SLICC) 2012 [18] SLE classification standard; 2) absence of other photosensitivity-related diseases, e.g., polymorphous light eruption, chronic dermatitis, solar urticaria, drug-induced phototoxicity, etc. [10]; 3) no history of cancer; 4) not pregnant at the time of diagnosis; 5) no diagnosis of other autoimmune diseases. Their demographic data, relevant clinical information, and serum test results – including SLE Disease Activity Index 2000 (SLEDAI-2K), anti-dsDNA antibodies, complement levels (C3, C4), erythrocyte sedimentation rate (ESR), total IgE (Immunoglobulin E), and whether lupus onset occurred on the face – were included as potential factors related to SLE activity. SLE disease activity levels were categorized according to the SLEDAI-2K scores as follows: 0–4 (stable/inactivity), 5–9 (mild activity), 10–14 (moderate activity), and ≥ 15 (severe activity) [22-24], with measurements taken on the day of hospitalization. Photosensitivity tests were conducted using the SUV-1000 sun simulator (1000W xenon short-arc lamp, Shanghai SIGMA High-tech Co., Ltd.). Reactions were assessed immediately and 24 hours after irradiation, recording UVB minimal erythema dose (UVB-MED) values. UVB-MED < 35 mJ/cm² was defined as UVB sensitivity. This study was approved by the Institutional Review Board of Huashan Hospital (HIRB), Fudan University (NO: 2013-015).

### Statistics

The comparation, association and correlation between SLEDAI-based disease activity, UVB-MED, and other factors were analyzed by employing Mann–Whitney U test, Kruskal–Wallis test, student t-test, one-way ANOVA, χ2 test, linear regression with Spearman correlation, locally weighted regression (Loess), B-spline, basis (polynominal) spline, RCS (restricted cubic spline) regression, polynomial regression, ROC (receiver operating characteristic) analysis, and logistic regression. *P*-values < 0.05 was interpreted as statistically significant. All statistical analyses were processed in R 3.6.3.

### Systemic collection of candidate UV-response genes/molecules

To collect candidate UV-response genes/molecules, we conducted a systematic literature search on PubMed, screening all English-language articles related to LE, UV, photosensitivity, photobiology, and photoprovocation published before June 20th, 2020. We used Medical Subject Headings (MeSH) and synonyms of key terms, combining them with the Boolean operators “OR” and “AND” for searches in titles and abstracts via Entrez. The search terms for UV-light included “ultraviolet rays”, “UV”, “UVB”, and “ultraviolet”. For lupus, we used “lupus erythematosus, systemic”, “SLE”, “lupus”, and “LE”. To capture photobiological aspects, we included “photosensitivity disorders/genetics”, “photobiology”, “photosensitivity”, and “photoprovocation”. Additionally, we reviewed references and citations from relevant articles to identify further studies (Fig. 2B, and Table S3A).

To maximize the inclusion of potential candidate genes, we adopted broader literature-selection criteria. We included all types of research papers – whether original articles, review articles, or correspondence letters – that provided information on the regulation of gene expression, protein levels, activity, or function in response to UV exposure (Fig. 2A, B). Two investigators independently screened the abstracts and full texts of each potentially eligible study. For all studies, the following data were extracted: gene symbol/name, biological regulation level/effects, tissue/cell source, LE type or other diseases, UV wavelength, study type (cell line/animal/human), and citation PMID (Table S3A). For some molecules reported in gene families, entire families of related genes (e.g., interferons and their receptors, MHC molecules, MMPs, collagens, and HSP70s) were included for further screening (Fig. 2A, B).

### Collection of UV-response genes/molecules from RNA-seq studies on UVB radiation in LE lesional/normal skin

To collect additional UV-response genes/molecules, we analyzed two clinical RNA-seq studies focusing on UVB radiation’s effect on LE lesional and/or normal skin. For the RNA-seq dataset GSE148535 from Skopelja-Gardner et al. [14], raw count data was downloaded from GEO supplementary files. The data was processed using both *limma/voom* and *Deseq2* program to calculate logarithm of fold change fold change (log_2_FC) and *P* values for multiple comparations: 24-hour radiation vs. non-radiation, 6-hour radiation vs. non-radiation, and 6- or 24-hour radiation vs. non-radiation groups. We then combined the differential gene expression analysis (DEA) results and retained only the highest absolute log_2_FC values for redundant genes by sorting and ranking them accordingly. Applying a cutoff standard of log_2_FC ≥ 1.5 and *P* value < 0.05, we compiled a list of 7,893 UVB-response differentially expressed genes (DEGs) for further analysis. Another RNA-seq dataset from Katayama et al. [11] focused on UVB radiation effects on normal, DLE, and SCLE human skin. We utilized their list of 411 UVB-response DEGs from a supplementary file for further analysis. There were 222 DEGs shared by Skopelja-Gardner and Katayama datasets. Notably, 200 of these genes did not overlap with the 296 UV-response candidate genes identified from the literature, and these are presented in Table S3B).

### Systemic collection, organization, quality control, and data processing of LE transcriptomic datasets

To screen and verify UV-response genes associated with LE disease activity, we systematically searched the GEO (Gene Expression Omnibus) database using MeSH terms, including “lupus erythematosus, systemic”, “lupus erythematosus, discoid”, “lupus erythematosus, cutaneous”, and “lupus nephritis”. We filtered the results by "*Homo sapiens*" (organism) and "Series" (Entry type) to ensure a comprehensive inclusion of relevant clinical transcriptomic microarray datasets. Inclusion and exclusion criteria were: 1) Inclusion of both human clinical LE and normal/control samples within a series; 2) no specific treatments (excluding conventional therapies) administered to patients or samples (if a series involves a specific medical therapy, only baseline samples were utilized); 3) for blood or skin tissue series, a minimum sample size of ≥ 5 in each LE or normal group; for other tissue/cell series, a minimum group size of ≥ 3; 4) All platforms must consist of single-channel arrays.

We then organized the selected series into working datasets (Fig. S2) by 1) splitting series that contained multiple tissue/cell types, and 2) merging series (or the previously split series) with the same platforms and tissue/cell types into new datasets using the batch-effect removal method *Combat* from *sva* R package. Regarding GSE65391, a large-sample series with two batches (R1 and R2), we maintained the independence of these batches in all subsequent analyses. For all datasets, the following data were recorded: series number, tissue/cell type, LE disease type, platform, and sample size (Table S4).

By June 20^th^, 2020, from 122 microarray series related to human clinical LE samples, 37 qualified series encompassing 5918 clinical samples were selected and then organized into 48 working datasets, which include eight CLE skin datasets (including one hair follicle dataset) and 40 SLE datasets. The SLE datasets comprise 12 whole blood datasets, five PBMC datasets, seven CD4 T cell datasets, one CD8 T cell dataset, five monocyte datasets, one macrophage datasets, four B cell datasets, one LDG (low density granulocyte) dataset, two neutrophil datasets, one platelet dataset, and one DC dataset (Fig. S2). Among these, eight SLE datasets included SLEDAI and other lupus activity-related clinical data, while three datasets provided therapeutic data and immunocyte flow cytometry data, which were utilized for further analysis. Detailed information about all the datasets is documented in table S4.

For microarray quality control and expression matrix extraction, we utilized R packages tailored to the different types of raw data from various platforms. For example, the *affy* package was employed to import and normalize Affymetrix U133 arrays, while the *oligo* package was used for processing Affymetrix whole transcriptomic arrays. Additionally, the *beadarray*, *limma*, or *vsn* packages were applied to process Illumina and Agilent microarrays. Missing values in any expression matrix were imputed using the KNN algorithm from the *Impute* package.

For gene annotation of microarray probes, we we prioritized downloading and utilizing the most recent annotation file from the official website of each platform. If the official annotation files were unavailable, the annotation files from GEO database were employed. In collecting clinical data for datasets and samples, we examined, combined, and filtered the data from supplementary files from GEO and related published article, as well as the phenotype data files generated by the *geoquery* package.

For DEA of microarrays, the *limma* package was employed to calculate log_2_FC, *P* value, and adjusted-*P* value for each gene. For GSE46907, which applied a microarray set composed of 2 arrays U133A and U133B, we combined the DEA results from both arrays and retained the results with the highest absolute log_2_FC for redundant genes after sorting them by absolute log_2_FC. Figures related to data quality control and distribution, group gene-expression differences, and DEA results are presented in Fig. S3.

### Lupus activity-correlated, UVB-response gene (UVBACG) identification

We presented log_2_FCs and adjusted *P*-values (or *P*-values for datasets with small sample sizes) of candidate UV-response genes from DEA with heatmaps (Fig. 2A and S4). For genes with multiple probes in an array, the probe with the highest log_2_FC was selected as the representative. We excluded IFNL4, HLA-DQB3, and IGKV3-20 from the candidate list, as most microarrays lacked corresponding probes.

Any gene in any tissue/cell type showing significantly differential expression between disease and normal samples was retained as a candidate LE UV-response DEG. These genes were then intersected with the DEG lists of Skopelja-Gardner and Katayama datasets, respectively, and merged into a single LE UVB-response DEG list (Fig. 2B).

Next, we performed both Spearman correlation analysis (between LE UVB-response DEGs and SLEDAI) and logistic regression (between LE UVB-response DEGs and SLEDAI activity groups) on eight datasets with available SLEDAI and/or other activity-related factor data. By filtering for Spearman correlation coefficient (r), odds ratio (OR) from logistic regression, and significant *P* values, any significantly correlated gene shared by at least two of the eight datasets was retained as a UVBACG. UVBACGs showing consistent changes in both DEA and Spearman correlation/logistic regression, as well as across all datasets, were ultimately collected as final consistent UVBACGs (Fig. 2B, C). Consistency was defined as a gene being either highly expressed in LE (compared to normal samples) and positively correlated with SLEDAI (and/or other activity-related factors) simultaneously, or showing the opposite pattern. The same filtration process was applied to the intersected DEGs between the Skopelja-Gardner and Katayama datasets to identify additional UVBACGs. The entire process flow is illustrated in Fig. 2B. A subset of UVBACGs that exhibited strong correlations with SLE activity and maintained high consistency across all datasets were considered key UVBACGs and selected for further analysis.

### Functional enrichment and protein-protein interaction analysis of UVBACGs

To illustrate the functions and pathways in SLE related to UVBACGs, we performed functional enrichment and protein-protein interaction (PPI) analysis by using Metascape [25]. For functional and pathway enrichment analysis, we included KEGG (functional sets, pathways, and structural complexes), GO (molecular functions, biological processes, cellular components), Reactome Gene Sets, BioCarta Gene Sets, PANTHER pathway, and WikiPathways with Metascape default settings: *P*-value < 0.01, minimum overlap count > 3, enrichment factor > 1.5. Regarding PPI analysis, we employed STRING, BioGrid, OmniPath, and InWeb_IM with Metascape default settings. Network figures generated by Metascape were further modified using Cytoscape (v3.9.0). We first carried out functional/pathway enrichment and PPI on the 39 UVBACGs.

Next, using the blood sample datasets GSE56391R1 and GSE56391R2, we filtered genes that were correlated with both UVBACGs and SLEDAI. Initially, we performed Spearman correlation with SLEDAI, retaining all genes with an absolute Spearman/Pearson r > 0.2 and *P* < 0.05. For each of these genes, Pearson correlation was then conducted with each UVBACG. For interferon-stimulated genes (ISGs) within the UVBACGs, we calculated an ISG score by summarizing the z-scores of each gene (details provided below). Genes with a Pearson r > 0.3 and *P* < 0.05 in correlation with the key UVBACGs or ISG score were retained and combined across both datasets. These genes were referred to as UVBACG/SLEDAI-correlated genes. As a result, each UVBACG had an associated list of UVBACG/SLEDAI-correlated genes, which were subsequently used for functional/pathway enrichment and PPI analysis. This approach helped identify the functional/pathway alterations and PPIs related to each UVBACG in SLE (results not shown). Finally, we defined key UVBACG/SLEDAI-correlated genes as those shared by more than four out of the seven key UVBACGs, yielding a final list of 273 genes. These UVBACG/SLEDAI-correlated genes, potentially representing key/common downstream targets triggered by UVBACGs in the immune system, were subjected to final functional/pathway enrichment and PPI analysis. The correlations between key UVBACGs and key UVBACG/SLEDAI-correlated genes were visualized through circular heatmaps (Fig. S6C).

### Interferon (IFN) signature score calculation

For IFN signature scores, we applied several scoring methods: Feng score [26], Skopelja-Gardner skin and blood score [14], Kirou IFNA and IFNG score [27], and Chaussasel M1.2, M3.4 and M5.12 scores [28]. These were calculated by summarizing the z-scores of each gene within the signature (1). We also applied the Petri score, which was calculated as the average of the z-scores (2). Since all probes corresponding to each signature gene were used (and different platforms contain different probes), only the Petri score was suitable for cross-platform comparisons across different microarray datasets.

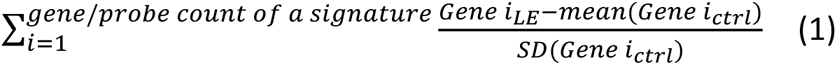

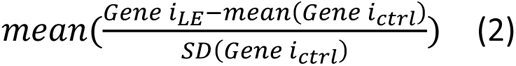

### Exploring cellular sources of UVBACGs in LE Patients

To demonstrate the correlation between cell abundance and key UVBACGs, and thereby identify the cell types in which the aberrant UVBACGs are localized, we utilized clinical flow cytometry data from datasets GSE49454, GSE65391-R1, and GSE65391-R2, alongside cell type abundance/enrichment estimations derived from gene expression matrices using bioinformatics tools. To enhance the reliability of these estimations, we employed three widely accepted tools that use different algorithms: xCell [29], ImmuCellAI [30], and CIBERSORT [31] following their default setting on both SLE blood samples (GSE49454, GSE65391R1, GSE65391R2) and CLE skin samples (GSE81071, GSE109248, GSE100093). Subsequently, we performed Spearman correlation analyses between cell enrichment/abundance and UVBACG expression. Any cell type showing a relatively high correlation (Spearman r > 0.2 with a *P* value < 0.05) with a UVBACG was considered a potential source of the UVBACG.

To further validate the expression of UVBACGs in the identified cell types, we examined immunohistochemistry staining of UVBACGs on skin tissue microslides from the Human Protein Atlas (HPA).

### Assessing Therapeutic Potential of UVBACGs in LE Patients

We applied three datasets with available therapeutic data (GSE49454, GSE65391R1, GSE65391R2) to evaluate the effects of regular anti-lupus treatments – hydroxychloroquine, corticosteroids (oral or intravenous), or immunosuppressive drugs (azathioprine, mycophenolate mofetil, methotrexate, cyclophosphamide, and cyclosporin A) – on regulating UVBACG expression. Here, we present the differential expression of key UVBACGs with or without these treatments, particularly across different SLEDAI levels.

To explore potential therapeutic targets further, we utilized the Comparative Toxicogenomics Database (CTD) [32, 33] for identifying possible chemical compounds, along with the SymMap database [34] and the Traditional Chinese Medicines Integrated Database (TCMID) [35, 36] to search for herbal ingredients. For each key UVBACG, we recorded potential chemicals and herbal ingredients from these databases. We performed intersection analyses within each database to identify candidates that might intervene in LE by regulating various aberrant UVBACGs. In the case of CTD, we focused on chemicals that negatively regulate ISGs, JUN, HLA-F, CAV1, and PTTG1, as well as those that positively regulate HOPX and RPL3, suggesting potential therapeutic effects. However, for SymMap and TCMID, precise information on how each herbal ingredient influences UVBACG expression (whether upregulated or downregulated) is lacking, indicating that further investigation is needed to clarify the regulatory effects of the identified herbal ingredients.

## Results

### Clinical characteristics

A total of 103 SLE patients with photosensitivity testing records from Department of Dermatology, Huashan Hospital between July 2011 and December 2017 were collected. Among them, there were 97 females and 6 males, with a mean age of 33.3 (14.0–72.0) years. Regarding disease activity, there are 21 (20.4%) stable (SLEDAI ≤ 4) cases and 82 active cases, the latter including 39 (37.9%) mildly active, 31 (30.1%) moderately active, and 12 (11.6%) severely active cases. As for UVB photosensitivity (UVB-MED), there were 16 patients (15.5%) not higher than 35 mJ/cm^2^ (UVB-sensitive reference value of our department) and 87 over it. Other demographic information, and clinical factors, including anti-dsDNA antibody, C3, C4, etc., are showed at Table 1.

**Table 1.**
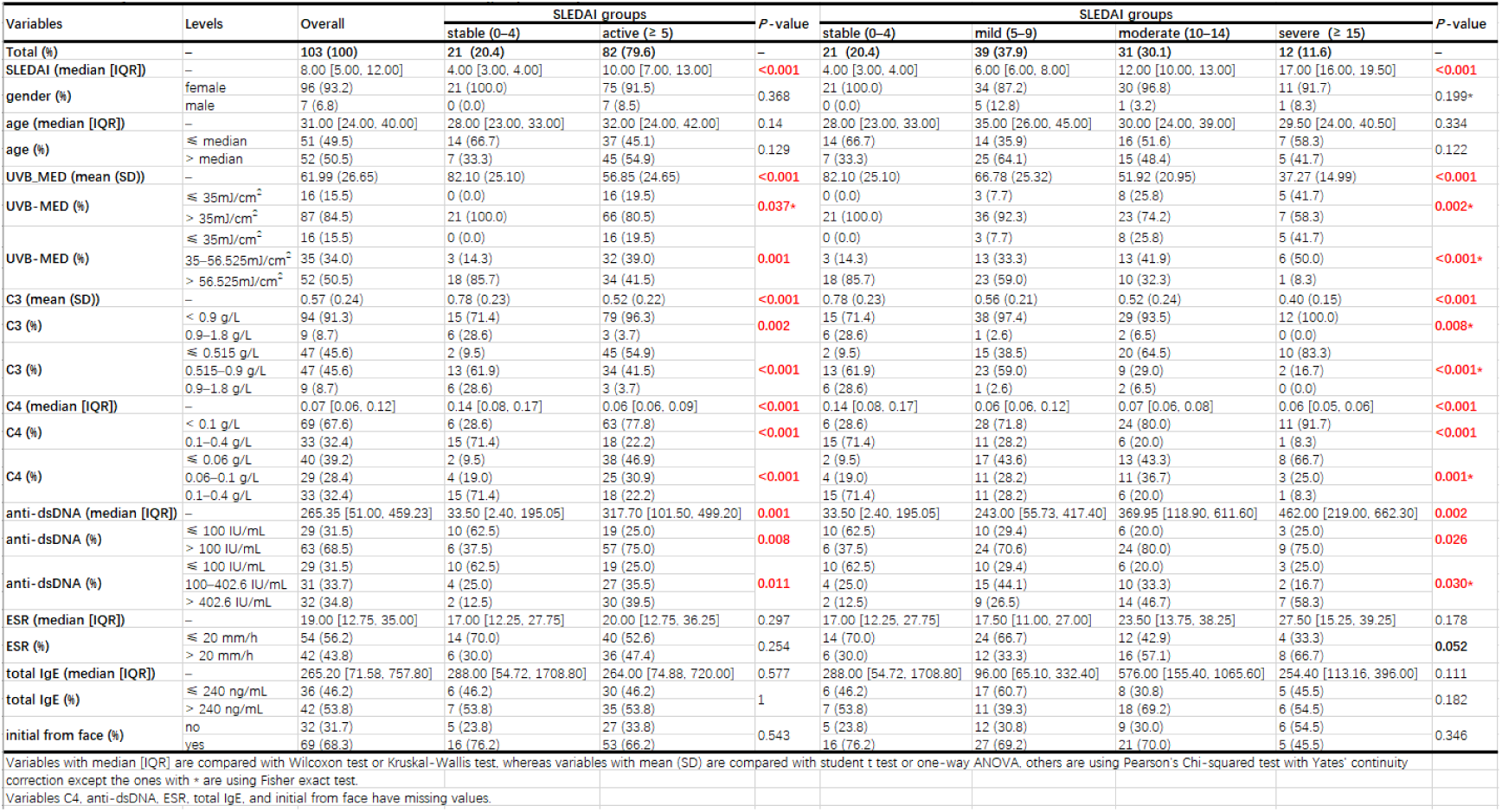
SLE activity-related factors and their associations with SLEDAI in SLE patients.

### UVB-MED, an indicator representing UVB photosensitivity, is significantly correlated with SLEDAI

To explore the relationship between SLEDAI and the clinical factors we collected, we first compared the value distribution of these factors across different SLEDAI groups, among which, UVB-MED, C3, C4, and dsDNA were significantly associated with SLEDAI (Table 1). Next, ROC analysis was performed to assess the ability of these factors to distinguish active SLE from stable cases. Notably, UVB-MED demonstrated a high AUC of 0.773 (*P* < 0.05, Fig. 1A), indicating its strong performance in discriminating SLE disease activity. The ROC Youden index J identified a UVB-MED value of 56.525 mJ/cm², which was extracted as a threshold (in addition to 35 mJ/cm²) for further analysis. Univariate logistic regression also demonstrated a strong correlation between UVB-MED and SLEDAI (Fig. 1B). Moreover, patients with a UVB-MED below 56.525 mJ/cm² had an 8.471-fold increased risk of SLE activity (*P* = 0.001) compared to those with values above this threshold (Table S1).

**Fig. 1.**
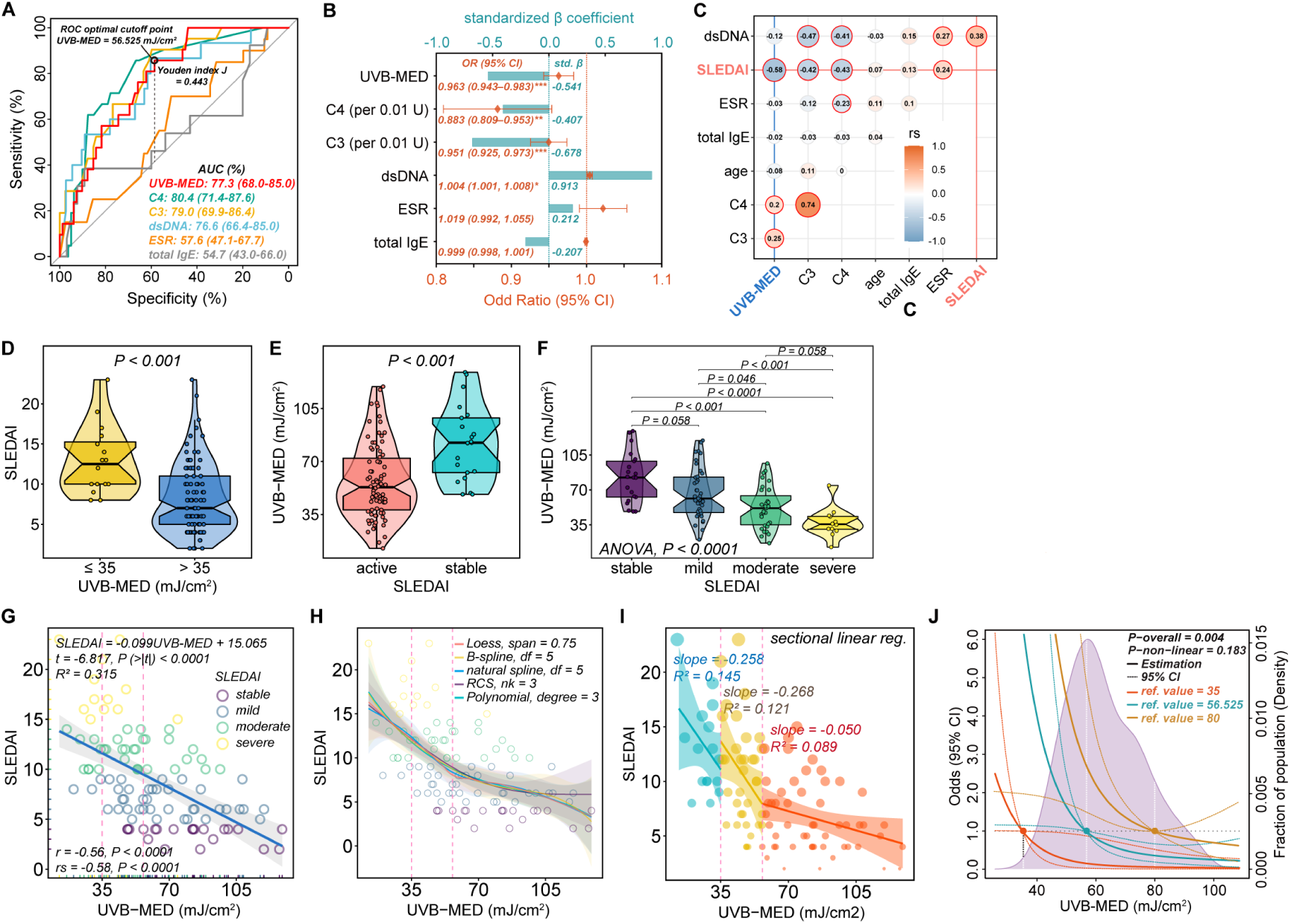
The association and correlation of UVB-MED and SLEDAI in SLE patients. **A)** ROC analyses for SLEDAI (active vs. stable) discrimination based on SLE activity-related factors. AUC, area under ROC curve. **B**) Univariate logistic regression for SLE activity-related factors (using exact values). Bar plots in cyan shows the standardized β coefficients (std. β) and forest plots in orange shows the odd ratios (OR) and 95% confidence intervals (CI) of each factor. **C**) Spearman correlation between SLEDAI, UVB-MED, and other factors. Colors and size of the circles indicate the spearman r. Red-bordered circle indicating P-values < 0.05. **D**) SLEDAI difference between UVB-MED groups (Wilcoxon test). **E–F**) UVB-MED difference between SLEDAI groups. E, student t-test; F, one-way ANOVA with post-hoc pairwise comparisons (holm method). **G**) Linear regression and Pearson/Spearman correlation between SLEDAI and UVB-MED. r, Pearson r; rs, Spearman r. **H**) Non-linear regression curves between SLEDAI and UVB-MED. Loess, locally weighted regression; B-spline, basis (polynominal) spline; RCS, restricted cubic spline; Polynomial, polynomial regression. The two dashed lines in G and H showing UVB-MED value: 35 and 56.525 mJ/cm^2^. **I**) Sectional linear regression between SLEDAI and UVB-MED. **J**) RCSs illustrating the non-linear correlation of univariate logistic odds and UVB-MED with different refence values (ref.). The dashed lines show 95% CI, and the density plot in purple shows the distribution of patients with different UVB-MED levels.

Next, multiple Spearman correlation analysis revealed that UVB-MED had the strongest correlation with SLEDAI (Fig. 1C). Notably, UVB-MED showed minimal correlation or association with other factors (Fig. 1C, Table S2). In multivariate logistic regression models, testing all combinations of UVB-MED, C3, C4, and dsDNA using continuous or ordinal data, UVB-MED consistently remained statistically significant across all models (Table S1), reinforcing its role as an independent risk factor for SLE activity. These findings clearly demonstrate and confirm that UVB-MED is an independent risk factor with a strong association and correlation to SLE disease activity, comparable to the well-established factors C3, C4, and dsDNA.

Focusing on UVB-MED, as shown in Fig. 1D, patients with lower UVB-MED (indicating higher UVB sensitivity) exhibited higher SLEDAI scores. Conversely, across different SLEDAI groups, those with more active disease (higher SLEDAI) had lower UVB-MED values (Fig. 1E, F). The relationship between UVB-MED and SLEDAI appeared to follow a dose-dependent-like pattern (Fig. 1F).

To illustrate quantitative relationship between UVB-MED and SLEDAI, we generated several regression curves, including linear, sectional linear, nonlinear, and spline regression analyses. Linear regression, along with Spearman and Pearson correlations, revealed a significant negative correlation (rs -0.58 and r = -0.56, *P* < 0.0001, Fig. 1G). Nonlinear regression curves further refined this relationship, showing a trend of gradually decreasing slope, indicating that as UVB-MED increases, the reduction in SLEDAI diminishes (Fig. 1H and S1). When we highlighted the ROC Youden index J-corresponding UVB-MED value (56.525 mJ/cm²) on the regression curves, it aligned with a turning point that divided the curve into two segments: one with steeper slopes and the other with more gradual slopes (Fig. 1H). The segment within the lower UVB-MED range exhibited a steeper decline, while the slope became more gradual and smoother at higher UVB-MED values. To confirm this trend, we exhibited a three-sectioned linear regression curve with segments corresponding to ≤ 35 mJ/cm², 35–56.525 mJ/cm², and > 56.525 mJ/cm². The first two segments displayed consistently steeper slopes, while the slope in the last segment was significantly reduced (Fig. 1I). By applying a restricted cubic spline to depict the UVB-MED value-dependent logistic odds for SLE activity levels (active vs. stable), we observed a trend similar to that of the previously established regression curves between UVB-MED and SLEDAI. As UVB-MED increased, the odds dropped exponentially, with the value of 56.525 mJ/cm² emerging as the turning point (Fig. 1J). Adjusting different reference values on the curves reinforced this trend. These results suggest that UVB-MED 56.525 mJ/cm² may serve as a more accurate threshold for UVB sensitivity grouping than 35 mJ/cm².

### Systematic literature mining of candidate UV-response genes

To explore the potential pathogenic mechanisms underlying UVB-triggered SLE inflammation, particularly focusing on the molecular targets involved, we conducted a systematic literature review and a comprehensive summary of related molecules and pathways. As a result, 296 genes, classified into 23 categories, were identified from 133 articles. These genes may be regulated at multiple biological process levels by UV radiation/exposure (Fig. 2A, B). The detailed list of these genes is provided in Table S3A. Two RNA-seq studies, Skopelja-Gardner et al. [14] and Katayama et al. [11], focusing on UVB radiation effects on LE lesional or healthy human skin, were utilized to identify UVB-induced differentially expressed genes (DEGs) (Fig. 2B). 7893 UVB-response DEGs were extracted, while 411 UVB-response DEGs were obtained from the Katayama dataset (Table S3B).

**Fig. 2.**
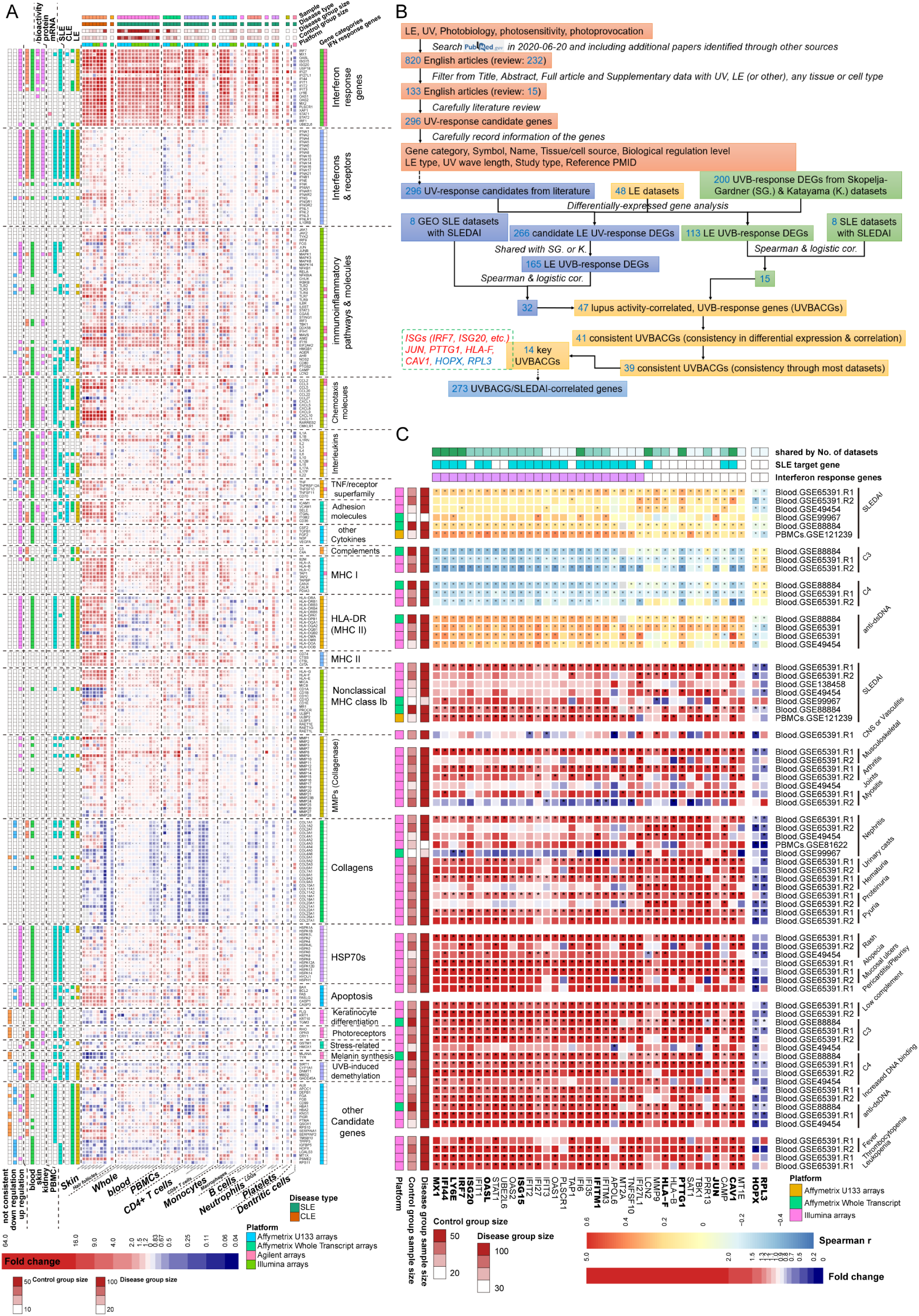
Systematically and comprehensively analyzing of UVB-response genes, as well as screening and identifying lupus activity-correlated, UVB-response genes (UVBACGs). **A**) A heat map showing differential expression of 296 UV-response candidates from systemic literature review in LE compared with normal samples of all kinds of tissues/cells. Color gradation showing log_2_FC, and asterisks showing P values < 0.05. **B**) A flowchart illustrating how UV-response candidate genes were collected and organized from systemic literature review, as well as how UVBACGs were identified. LE, lupus erythematosus; DEGs, differentially expressed genes. **C**) A heat map showing Spearman correlation r (for continuous factors) and logistic odd ratios (for ordinal factors) between the 39 consistent UVBACG expression with SLE activity-related factors in datasets. Orange-light blue color gradation showing spearman r, while red-dark blue color gradation showing logistic odd ratios, which were labeled with asterisks showing P values < 0.05. The 14 key UVBACGs are showing in bold.

### Systematic collection/organization of LE-related microarray datasets and identification of lupus activity-correlated, UVB-response genes (UVBACGs)

We conducted a systematic search of the GEO (Gene Expression Omnibus) database and identified 37 qualified series comprising 5,918 clinical samples, which were organized into 48 datasets (eight CLE skin datasets and 40 SLE blood/blood cell datasets, Fig. 2B and Fig. S2). Detailed information on all datasets is provided in Table S4.

Then, a comprehensively mining and screening of UVBACGs were conducted using these clinical transcriptomic datasets. We first performed differential expression analysis on all 296 UV-response candidates identified from the literature across the 48 LE datasets (Fig. S3). A heatmap overview revealed that most genes were significantly altered between LE and normal skin tissues, with deeper red or blue colors representing greater increases or decreases in expression. While, a relatively smaller number of genes were altered between patients and healthy donors in blood samples or in various types of blood cells. Remarkably, the most consistent DEGs between skin and blood samples were the interferon (IFN) I response/stimulated genes (ISGs) (Fig. 2A, see the trend from left to right). By excluding genes that were not differentially expressed between LE and normal samples, we identified and retained 266 DEGs, which we designate as candidate LE UV-response DEGs (Fig. 2A, B).

By intersecting these 266 genes with UVB-response DEGs from the Skopelja-Gardner and the Katayama datasets, we identified 165 LE UVB-response DEGs. These genes are likely significant in UVB stimulation of the skin and may also be retained by immune cells when disseminating into the bloodstream and other organs. Finally, we identified 32 UVBACGs that showed significant logistic and Spearman correlations with SLEDAI (Fig. 2B, C). These genes are not only closely associated with UVB-induced LE immune responses but also play key roles in LE activity.

To expand the list of UVBACGs, we also intersected the DEG lists from the Skopelja-Gardner and Katayama datasets. As a result, we identified an additional 200 candidate UVB-response DEGs shared between the two datasets (Table S3B, Fig. S4), distinct from the abovementioned 296 candidates. Following a similar identification process, we ultimately identified 15 additional UVBACGs (Fig. 2B, C).

After excluding six genes that were either highly expressed in LE samples but negatively correlated with SLEDAI, or vice versa, along with two additional genes showing significant inconsistencies across datasets, we ultimately identified 39 consistent UVBACGs from the initial 47 UVBACGs. Among these 39 genes, 14 exhibited the strongest correlation with SLE activity and maintained high consistency across all datasets. This subset included eight ISGs – IRF7, ISG20, ISG15, IFI44, IFITM1, MX1, LY6E, and OASL – which were positively correlated with SLE activity. Additionally, six non-ISGs were identified: JUN, PTTG1, HLA-F, and CAV1, also positively correlated with SLE activity, while HOPX and RPL3 showed a negative correlation with SLE activity (Fig. 2B-C). These 14 UVBACGs were considered key UVBACGs and selected for further analysis.

### Detecting difference in 1) UVBACG expression among healthy, patient non-lesional, and lesional skin tissues, as well as 2) UVBACG skin expression across CLE subtypes

As shown in Fig. S5, lesional LE skin follicles exhibited significant alterations in the expression of ISGs and HLA-F compared to non-lesional sites or healthy donors. Notably, LE non-lesional follicles also displayed elevated expression of ISGs and HLA-F compared to healthy donors, suggesting that UVBACG dysregulation and UVB sensitivity may already be present in the non-lesional skin of LE patients. However, no consistently differential UVBACG expression patterns were observed among the different CLE subtypes.

### Functional enrichment and protein-protein interaction analysis of UVBACGs

Functional enrichment and protein-protein interaction analysis of the 39 UVBACGs revealed significant enrichment in pathways related to interferon signaling (particularly IFN I), viral response, and immune system activation (Fig. S6A, B). Since our methodology identified UVBACGs as genes activated by UVB in LE skin tissues, it’s possible that we may have missed key molecules or pathways that become dysregulated in blood cells as a downstream effect of UVBACG dysregulation initiated from the skin. To address this, we screened and identified 273 genes from blood sample datasets that were correlated with both UVBACGs and SLEDAI (UVBACG/SLEDAI-correlated genes, Fig. 2B and S6C). In addition to their roles in interferon signaling and the adaptive immune response – particularly against viruses, these genes also showed involvement in cell cycle regulation, cell death and senescence, DNA replication, and eukaryotic translation elongation (Fig. 3). This suggests the mobilization and amplification of systemic immune responses, supporting the notion that UVBACGs are initially activated in the skin and subsequently trigger systemic pathogenic expansion after immune cells infiltrate the circulation and other body systems. Analysis of IFN signature scores between LE and control samples revealed aberrant IFN activity across tissue and cell types –including skin, blood, PBMCs, neutrophils, T/B cells, monocytes/macrophages, and platelets – with the largest differences observed in the IFNα signature, followed by IFNβ, and then IFNγ (Fig. S7).

**Fig. 3.**
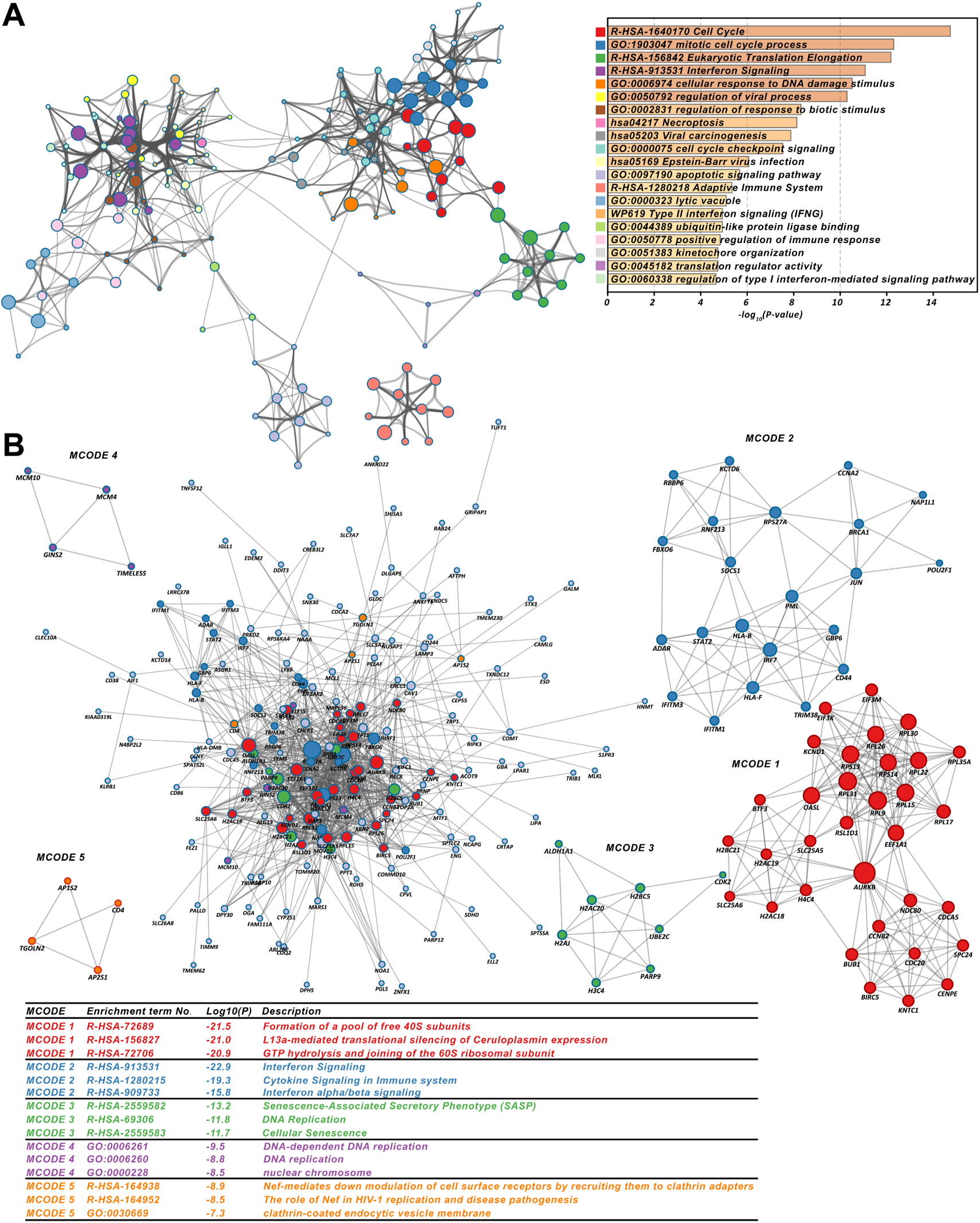
Functional enrichment (**A**) and protein-protein interaction (**B**) analyses of 273 key UVBACG/SLEDAI-correlated genes.

### Exploring cellular sources of UVBACGs in LE patients

To pinpoint the specific cell types where UVBACGs are likely to play a role, we conducted Spearman correlation analysis between UVBACG expression and cellular abundance data obtained through flow cytometry and computational estimation methods (xCell, CIBERSORT, and ImmuCellAI). Cell types with a Spearman correlation coefficient (r) > 0.2 and *P*-value < 0.05 were considered potential sources of UVBACGs. We selectively presented the results for IRF7 (as a representative ISG), JUN, and PTTG1 (Fig. 4A). A general summary of the cellular source estimation for all UVBACGs is illustrated in Fig. 4B. Most of the key UVBACGs were aberrantly expressed in the skin cells, with significant correlations observed between these genes and epidermal keratinocytes, as well as skin-resident immune cells – such as dendritic cells, neutrophils, CD4, CD8, and B cells (Fig. 4B). This indicates that the skin is the primary site for inflammation outbreaks. Moreover, all key UVBACGs were correlated with CD4, CD8, and B cells in both the skin and blood, as well as with γδT cells in the blood, suggesting that UVBACGs are retained within these cells as they migrate between the skin and bloodstream.

**Fig. 4.**
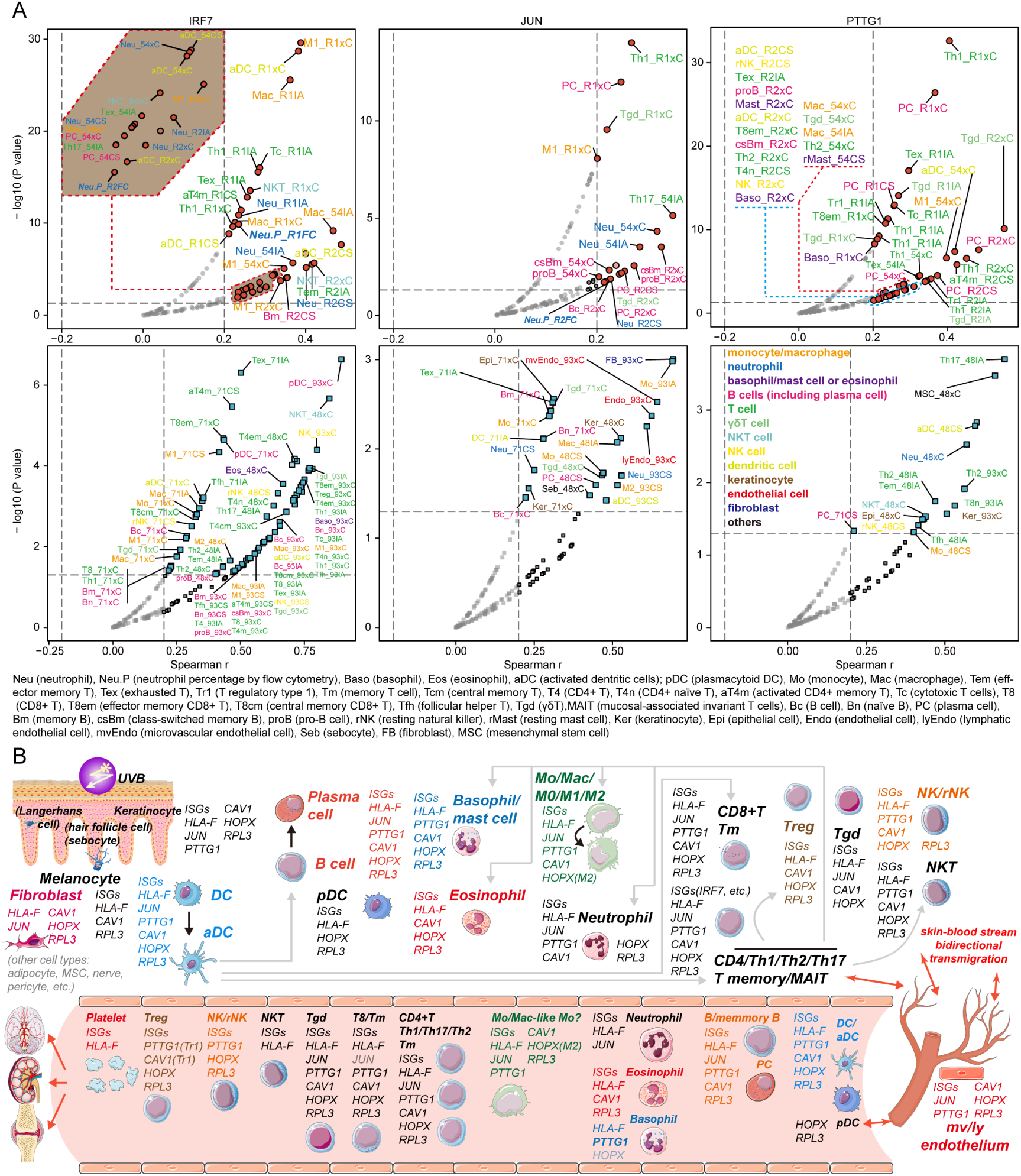
Exploring cellular sources of key UVBACGs in LE patients. **A**) Spearman correlations between key UVBACG expressions and cell type abundance based on computational estimation methods and flow cytometry. Upper panel: SLE blood samples, lower panel: CLE skin samples. The cell type with higher Spearman r and - log10 (P value) has the more likelihood the source of the gene. Abbreviation Labels showing cell type, datasets, and methods (e.g., M1_R1xC means xCell estimation of M1 macrophage abundance in GSE65391-R1). xC, CS, IA, and FC are referred to xCell, CIBERSORT, ImmuCellAI, and flow cytometry; 54, R1, R2, 71, 93, 48 are referred to GSE49454, GSE65391-R1, GSE65391-R2, GSE81071, GSE100093, GSE109248. **B**) Schematic model showing the potential cellular sources of UVBACGs in skin and blood, as well as our assumption on the role of UVB on LE activity. Cartoon images were adopted and modified from Servier Medical Art.

From the perspective of UVBACGs, ISGs were shared by all cell types, followed by HLA-F, HOPX, and RPL3. Some UVBACGs showed increased correlations from the skin to the blood cells, such as IRF7 in neutrophils, JUN in neutrophils and B cells, PTTG1 in B and γδT cells, as well as RPL3 in macrophages and γδT cells (Fig. 4A, B). The expression of UVBACGs in skin cells was further supported by immunohistochemistry staining of skin tissue slides from the Human Protein Atlas (HPA) (Fig. S8, S9).

### Assessing therapeutic potential of UVBACGs in LE patients

Using three SLE datasets GSE65391-R1, GSE65391-R2, and GSE49454, we demonstrated the potential impact of regular therapeutic drugs – hydroxychloroquine (HCQ), corticosteroids (CS), and immunosuppressive agents (IS, e.g., mycophenolate mofetil and cyclophosphamide) – on the expression of UVBACGs in SLE patients, particularly across different SLEDAI levels. The results for IRF7 (as a representative of ISGs), JUN, and PTTG1 are shown in Fig. 5. Consequently, HCQ, CS, and IS exhibited downregulatory effects on UVBACGs, with the effects being more pronounced in active patients compared to stable ones. This observation underscores the therapeutic benefits of conventional drugs for LE patients, particularly those with active disease, likely due to their role in correcting UVBACG dysregulation in LE. Notably, HCQ demonstrated the most significant and consistent decrease in the expression of IRF7 and PTTG1.

**Fig. 5.**
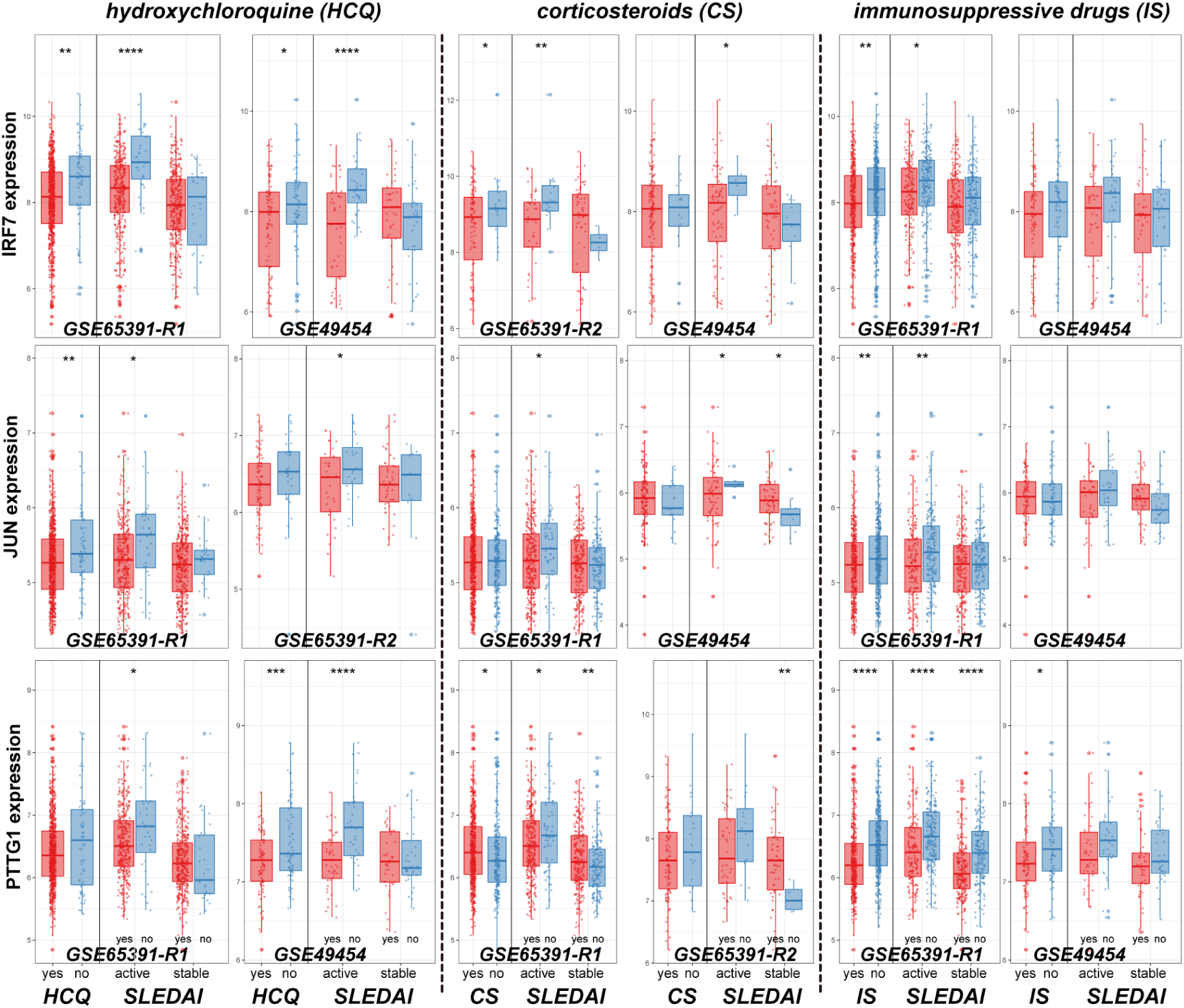
Hydroxychloroquine (HCQ), corticosteroids (CS), and immunosuppressive drugs (IS) may exert their therapeutic effects in SLE by regulating key UVBACGs. Boxplots showing the expression difference of three representative UVBACGs in different SLEDAI levels, with or without using HCQ, CS, or IS.

For further exploration of potential therapeutic drugs, we applied the SymMap database, TCMID (Traditional Chinese Medicine Integrated Database), and CTD (Comparative Toxicogenomics Database) to identify potential compounds (e.g., immunosuppressive, anti-cancer agents, etc.) and herbal ingredients that can simultaneously regulate a variety of aberrant UVBACGs (Fig. S10).

## Discussion

The subjectivity nature of photosensitivity, primarily due to the broad definition of photosensitivity in the ACR criteria [16], introduces uncertainty regarding the precise relationship between photosensitivity and LE. Consequently, the reported incidence of photosensitivity varies significantly within each type of LE; for example, subacute cutaneous lupus (SCLE) ranges from 27% to 100%, discoid lupus erythematosus (DLE) ranges from 25% to 90%, and SLE ranges from 6% to 54%, demonstrating considerable variation [2, 12, 20, 37]. Additionally, further studies have confirmed significant discrepancies regarding photosensitivity between patients’ complaint and photosensitivity tests [20]. Therefore, relying solely on patient complaint/history or physician observation for assessment is so subjective that the later updated criteria SLICC [18] and EULAR [19] gradually removed the item related to photosensitivity. The incorporation of a standardized laboratory test, such as phototesting, holds significant value in providing an objective measurement of photosensitivity. However, phototesting for photosensitivity has not yet been widely adopted as a routine clinical assessment, and no targeted intervention specifically addresses photosensitivity-related LE pathology. This may be attributed to: 1) the lack of accessible and practical photosensitivity indicators from UVB phototesting that are clinically validated and correlated with LE progression and activity, and 2) the absence of robust, systematic, and comprehensive evidence from experimental and clinical studies supporting UVB’s role in LE progression, particularly in relation to increasing photosensitivity.

This study aims to address these two challenges by providing potential solutions. For the former, we employed UVB-MED as an objective indicator of laboratory phototest to represent UVB photosensitivity. Dourmishev et al. found that LE patients had significantly reduced UVB-MED compared to healthy individuals, with increased UVB sensitivity more frequently observed in SLE and DLE [38]. Similarly, Reefman et al. reported that SLE patients had a significantly decreased UVB-MED [39]. Building and expanding on this, we demonstrated a close negative correlation between UVB-MED and SLE activity, revealing that exacerbation of LE is accompanied by increased UVB photosensitivity. The employment of a quantitative score allows us to quantify and grade UVB sensitivity, rather than simply dichotomizing patients into sensitive or not. Additionally, UVB-MED is a relatively independent risk factor for disease activity compared to other SLE serum markers (C3, C4, dsDNA, etc.). Therefore, UVB-MED has the potential to serve as a diagnostic indicator for high UVB sensitivity SLE, or as a supplementary component to existing SLE activity assessment systems.

Further, we generated a potential novel UVB-MED reference value 56.525 mJ/cm^2^ to define the UVB photosensitivity level, which is worth of verifying by further studies. Patients with a UVB-MED below this value can be more susceptible to UVB-induced SLE exacerbation and therefore require special attention and increased follow-up to prevent disease progression triggered by exposure to sunlight or other sources of high-dose UVB.

For the latter, given that current published experimental studies using cell and animal models are insufficient to fully reflect human pathological conditions, and that clinical studies often lack systematic and comprehensive methodologies [11, 14, 40], we systematically reviewed, organized, summarized, and verified most candidate genes from the literature. By employing nearly all available LE transcriptomic datasets (5918 clinical samples, 48 datasets), we performed comprehensive analyses to screen, filter, and identify lupus activity-correlated, UVB-response genes (UVBACGs). Furthermore, we explored their potential cellular sources, pathological roles, and possible therapeutic interventions. These clinical samples/datasets offer significant advantages: they represent real-world clinical objects exposed to sunlight, which enable efficient discovery, evaluation, and verification of UVBACGs from candidate genes identified in literature from heterogeneous studies. Therefore, our mining process ensured that the UVBACGs we identified meet key criteria: 1) relevant to real-world clinical scenarios, 2) dysregulation by UVB irradiation experiments, 3) aberrant expression in LE samples with a correlation to disease activity, and 4) validation by existing literature.

We incorporated both CLE and SLE into some analysis, due to the facts that they are closely related – 1) progression of CLE to SLE, 2) a notable proportion of SLE patients developing CLE-specific lesions, and 3) the coexistence of SLE and CLE [2, 3, 12] – that UV sensitivity and skin UV radiation-associated pathology are common features of multiple LE subtypes. Here, we found that non-lesional skin of DLE patients might already exhibit UVBACG dysregulation and UVB photosensitivity. This is supported by several other studies which demonstrated that non-lesional keratinocytes express ISGs more than those in healthy controls [41, 42]. These results imply that CLE is an overall skin disease, with inflammation not confined to the rash areas but instead in a state of dynamic spread. Plus, during follow-up, CLE patients who eventually develop SLE gradually exhibit systemic manifestations before fulfilling the criteria for SLE [6]. Notably, sun exposure can trigger systemic symptoms from CLE [12]. Therefore, it is highly likely that the expansion and exacerbation of inflammation promoted by UV from the skin to the system links CLE and SLE together. Our comprehensive study helps uncover the overall pathogenesis of how CLE progresses to SLE and how SLE manifests with CLE.

First, highlights UV-provoked skin as the “primary crime scene” where injury, autoimmune, and both innate and adaptive immunopathology unfold. We found that the majority of UV response candidate genes were significantly altered in LE skin tissues compared to normal skin, while fewer genes remained altered in the bloodstream or various blood cell types. Most UVBACG expression was further confirmed to correlate with various inflammatory cells in the skin, including resident keratinocytes, fibroblasts, and immune cells such as neutrophils, eosinophils, basophils, macrophages, T cells, B cells, γδT cells, NK cells, NKT cells, and dendritic cells. This is supported by literature that UVB acts as a trigger for a malfunctional immune state of skin through multiple mechanisms. 1) It is well-established that UVB induces damage and apoptosis in epidermal cells, thereby compromising the skin barrier, leading to microbial invasion and/or disrupting the pre-existing immune balance of the skin [4]. Patra, et al. reported that UVB induces a pro-inflammatory environment in the skin when microbes are present, but promotes an immunosuppressive state when microbes are absent, indicating that UVB damage to the epidermal immune barrier facilitates microbial invasion [43]. 2) Skin cell damage/injury itself also initiates injury-inflammation response, e.g., UVB-induced SLE apoptotic keratinocytes release high-mobility group protein B1 (HMGB1) and other inflammatory initiators, such as ATP. The latter is an early cellular stress/death signal, which can further activate human skin-resident T cells and mouse dendritic epidermal γδT cells [44, 45]. 3) Translocation of autoantigens (Ro/SSA, La/SSB, etc.) from the nucleus/cytoplasm to the apoptotic keratinocyte surface powered by UVB is another important mechanism [42]. UVB can promote LE by relocating SSA/Ro to the surface of basal keratinocytes [46, 47]. Increased autoantigens and apoptotic cells exceed phagocytes’ clearance capacity, which significantly enhances the opportunity for antigen presentation to immune cells [48]. 4) UVB radiation can also promote human endogenous retroviruses dsRNA expression thus activating dsRNA-sensing antiviral RIG-I/MDA5/IRF7 pathway, which induces IFN-I production, potentially serving as a mechanism for the inflammatory response and skin lesions observed in SLE/DLE [49].

Then, UVB-induced immune dysregulation worsens and expands uncontrollable skin inflammation, both innate and adaptive immunity involved, manifesting as photosensitivity in the skin. 1) The ongoing disruption of skin immune homeostasis (barrier breakdown, cell injury, autoantigen translocation, etc.) recruits numerous inflammatory immune cells, initiating a forward feedback loop that amplifies autoimmunity. For instance, UVB-induced skin lesions in SLE increase IFN-I in the epidermis and dermal endothelium, thus upregulating endothelial E-selectin and ICAM-1, leading to T cell and monocytes/macrophage recruitment. Elevated IFN-I further triggers stronger pro-inflammatory and chemotactic signals, amplifying skin inflammation [50]. 2) UVB can directly activate immune cell functions. UVB exposure can induce DNA hypomethylation in CD4+ T cells by overexpressing GADD45A [51], through the AhR-SIRT1-DNMT1 axis [52], and by inhibiting DNMT3A [53], among other mechanisms. This can result in the overexpression of genes such as CD11a, CD70, perforin, and CD40 ligand, leading to T cell autoreactivity and proliferation, monocyte/macrophage killing, and excessive B cell stimulation with IgG overproduction [51]. Additionally, UVB induces IL-6 production in the macrophages/monocytes [54] and alters the cytotoxicity of NK cells from SLE patients [55], as well as induces ROS-dependent neutrophil extracellular trap (NET) formation in vitro [56].

Second, our study suggests that skin inflammation extends into the bloodstream and spreads throughout the body, contributing to systemic autoimmunity. This explains how CLE progresses to SLE and how UVB exposure exacerbates disease activity in SLE, manifesting as systemic photosensitivity. Sun-exposed skin of patients with sun-induced systemic symptoms (compared to those in whom sun exposure does not trigger systemic symptoms) shows an increased presence of myeloid dendritic cells and T cells, suggesting that LE systemic reactions are linked to, and possibly driven by, the expansion of skin inflammation [12]. There are two mechanisms involved: 1) Proinflammatory factors secreted by skin inflammatory cells can disseminate into the bloodstream and throughout the body. Our study revealed that the difference in ISG expression between LE and healthy individuals is more pronounced in skin samples compared to blood/blood cell samples. Recent reports demonstrated that UVB-induced IFN signatures from the blood and kidney, can originate from skin IFN-I dissemination from skin [13, 14]. Skin pDC and keratinocytes might be the main sources of IFN-I in SLE [14, 57, 58]. Other proinflammatory cytokines, such as IL-17 and IL-22, can be released from skin lymphocytes into the circulation, potentially contributing to systemic inflammation [59]. 2) Skin inflammatory cells can directly disseminate to the circulation and other organs, maintaining aberrant UVBACG expressions. These aberrant expressions can further promote autoimmune functions by dysregulating additional immunopathways or manipulating other cells, thereby triggering systemic inflammation. We illustrated that all the key UVBACGs are correlated with CD4, CD8, and B cells in both the skin and blood, as well as with γδT cells in the blood, suggesting that UVBACGs are maintained within these cells as they migrate from skin to the bloodstream. Some UVBACGs showed increased correlation from the skin to the bloodstream. As such, IRF7, a representative of ISGs, exhibited a marked increase in correlation in neutrophils. Additionally, JUN in both neutrophils and B cells, PTTG1 in B cells and γδT cells, as well as RPL3 in macrophages and γδT cells were elevated. Supporting our results, recent investigations confirmed that neutrophils, crucial in LE inflammation, not only recruit to UV-exposed skin, but also spread systemically. As a response of UV-induced IFN-I, the IL-17A/G-CSF axis, drives neutrophilia and homing to tissues [13, 14]. In the kidney, neutrophils exhibit a proinflammatory profile, including increased ROS production, release of tissue-damaging proteases such as elastase and MMP8 (as observed also in our study), and NETs formation. A subset of neutrophils has been proved to extravasate from UV light-exposed skin tissue through reverse transmigration, contribute to the kidney IFN-I signature and initiate the inflammation and injury of kidney [13].

Among all the UVB-response genes in our study, ISGs are the most consistent genes found in both skin and blood across various cell types. Additionally, UVBACGs displayed significant features of interferon signaling (especially IFN-I), response to virus infection, and immune system activation. Indeed, a solitary UVB exposure is sufficient to induce a robust IFN-I response in the skin of both mice and humans [14]. UV irradiation-induced DNA damage, keratinocyte apoptosis, immune barrier compromise, autoantigen translocation, and promotion of endogenous retroviruses dsRNA have all been reported to initiate and enhance IFN-I responses in LE [11, 48, 49, 60, 61]. Moreover, IFN-I can stimulate nearly all immune cells [14, 62, 63]. Notably, IFN-I can induce SLE monocyte differentiation into DCs with potent antigen presentation capabilities [64]. Besides interferon signaling and the adaptive immune response (particularly against viruses), functional enrichment of UVBACG/SLEDAI-correlated genes demonstrated phenotypes related to cell cycle, DNA replication, and eukaryotic translation elongation. This implies the mobilization and amplification of systemic immune responses after UVBACGs are initially activated by UVB in the skin.

Other UVBACGs, including six non-ISGs (JUN, PTTG1, HLA-F, CAV1, HOPX, and RPL3), though less studied, are worth further investigation for their potential involvement in the mechanism by which photosensitivity forms from skin to the system. We have detailed the roles of these genes in LE and immunity, as well as their interactions with UV, in Supplementary Discussion.

The association between UVBACG expression and regular LE treatments (hydroxychloroquine, corticosteroids, and immunosuppressive medications) suggest that these treatments, particularly for active SLE, may exert their effects by regulating key UVBACGs. This is supported by literature showing that the regular treatments reduce levels of ISGs in patients [65, 66]. Further, pre-treatment of mice with hydroxychloroquine significantly reduced IFN-I scores in both skin and blood after UVB exposure [14]. However, consider the side effects from long-usage of the regular medications, exploration of novel targeted therapy to UVBACGs and their related pathways is of great necessity. Moreover, we emphasize the use of UVBACG-targeted topical agents, along with sunscreen, considering that the skin is the “primary crime scene” of LE. For more insights into our views on how to target UVBACGs for treating LE, please refer to the details in Supplementary Discussion.

To summarize the key points of the two major parts of this study (Fig. S11). UVB, as a crucial factor in LE diseases, can trigger the onset of LE, advance CLE to SLE, promote disease activity, and manifest SLE with CLE lesions. When enough area of the skin is exposed to a sufficient dose/time of UVB radiation, factors such as cell damage and autoantigen exposure, etc., drive the outbreak of skin inflammation, followed by autoimmune expansion, and systemic dissemination, with IFN-I playing a predominant role [13, 14, 67, 68]. Regarding photosensitivity of LE patients, to note, they have been carrying an existing autoimmune state (a pre-lesional state), where cellular UVBACGs, particularly ISGs, have already deviated from normal levels, lowering the threshold for the onset of skin and systemic inflammation [42]. The more active the autoimmune cellular state in a patient, the lower the threshold for inflammation, indicating heightened photosensitivity. When the minimum dose of UVB exposure sufficient to trigger skin inflammation is measured, known as UVB-MED, it can be used to objectively quantify photosensitivity.

## Supporting information

Supplementary Discussion

## Supporting information

### Supplementary discussion

See online Supplementary Discussion.pdf

### Supplementary table legends

**Table S1.** Univariate and multivariate logistic regression analyses for SLE activity status (active vs. stable) using multiple disease activity risk factors, including UVB-MED.

**Table S2.** The associations of SLEDAI and SLE activity-related factors with different UVB-MED levels in SLE patients.

**Table S3.** UV-response candidate gene lists. A) List of 296 UV-response candidate genes and their detailed information from systemic literature review. B) List of additional 200 UVB-response candidate genes from the studies by Skopelja-Gardner et al. and Katayama et al., complementing the 296-gene list in part A.

**Table S4.** List of 48 LE GEO datasets and their detailed information.

### Supplementary figure legends

**Fig. S1. Non-linear regressions between SLEDAI and UVB-MED.** Illustration of non-linear regressions employing gradually varied parameters for each regression method: Loess (locally weighted regression), B-spline (basis polynomial spline), RCS (restricted cubic spline), and Polynomial regression. The dashed lines in all figures indicate the thresholds of 35 and 56.525 mJ/cm2.

**Fig. S2. Flow chart summarizing the structure and content of the present study.** This comprehensive flow chart illustrates the overall structure and content of the study. Abbreviations: LE (lupus erythematosus), DE (differential expression), DEGs (differentially expressed genes).

**Fig. S3. Boxplots, PCA plots, and volcano plots illustrate the data distribution, group differences, and differentially expressed genes across the 48 GEO SLE datasets included in this study.**

**Fig. S4. A heatmap displaying differential gene expression (fold change) for LE samples compared to normal samples across the 200 UVB-response candidate DEGs identified by Skopelja-Gardner et al. and Katayama et al., in addition to the 296-gene list.**

**Fig. S5. Differential expression of UVBACGs in different CLE subtypes (A-C), as well as between healthy, patient non-lesional, and lesional skin follicle tissues (D).** * *P* < 0.05, ** *P* < 0.01, *** *P* < 0.001, **** *P* < 0.0001.

**Fig. S6. A-B**) Functional enrichment and protein-protein interaction analysis of 39 consistent UVBACGs showed significant features of interferon signaling (especially IFN I), response to virus infection, and activation of type I immunity related intracellular pathways. **C**) Circular heat maps showing the expression correlation of key UVBACG/SLEDAI-correlated genes with key UVBACGs (Pearson) and SLEDAI (Spearman), using GSE65391-R1 and R2 datasets.

**Fig. S7.** Comparisons of IFN signature scores between LE samples (CLE in green and SLE in purple) and control samples across all datasets, based on published IFN signature genes [Feng score, Skopelja-Gardner skin and blood score, Kirou IFNA (IFNα) and IFNG (IFNγ) score, Chaussasel M1.2 (IFNα), M3.4 (IFNβ) and M5.12 (IFNγ) scores]. The results indicate that most tissue/cell samples, including skin, blood, PBMCs, T/B cells, monocytes/macrophages, and platelets, exhibit the highest differences in the IFNα signature, followed by IFNβ, and then IFNγ.

**Fig. S8.** Potential cellular source evidence of UVBACG expression by immunohistochemistry staining on skin tissue microslides from Human Protein Atlas (HPA) (https://www.proteinatlas.org/).

**Fig. S9.** Other valuable immunohistochemistry skin tissue microslides from HPA which can support cellular source evidence of UVBACGs. We suggest readers who are interested at these slides to check them in the website of HPA (https://www.proteinatlas.org/).

**Fig. S10. Applying TCMID (Traditional Chinese Medicine Integrated Database, A), SymMap database (B), and CTD (Comparative Toxicogenomics Database, C) for identification of possible chemicals and herbal ingredients that might ameliorate disease activity by simultaneously regulating a variety of aberrant UVBACGs.** Green, anti-inflammation chemicals; red, immunosuppressive or anti-tumor chemicals; blue, sex hormone-related chemicals. ISG: interferon-stimulated genes. The horizontal (green, orange, and blue) columns display the number of substances in the database that can regulate the corresponding UVBACG, while the vertical (black) columns show the number of substances in the database that can simultaneously regulate several UVBACGs, as indicated by the black dots in the diagram below.

**Fig. S11. An illustrated summary of the key findings from this study.** UVB, as a crucial factor in LE diseases, can trigger the onset of LE, advance CLE to SLE, promote disease activity, and manifest SLE with CLE lesions. When enough area of the skin is exposed to a sufficient dose/time of UVB radiation, factors such as cell damage and autoantigen exposure, etc., drive the outbreak of skin inflammation, followed by autoimmune expansion, and systemic dissemination, with IFN-I playing a predominant role. Regarding photosensitivity of LE patients, to note, they have been carrying an existing autoimmune state (a pre-lesional state), where cellular UVBACGs, particularly ISGs, have already deviated from normal levels, lowering the threshold for the onset of skin and systemic inflammation. The more active the autoimmune cellular state in a patient, the lower the threshold for inflammation, indicating heightened photosensitivity. When the minimum dose of UVB exposure sufficient to trigger skin inflammation is measured, known as UVB-MED, it can be used to objectively quantify photosensitivity.

