## Supplementary Discussion for "Exploring the Correlation Between UVB Sensitivity and SLE Activity: Insights into UVB-Driven Pathogenesis in Lupus Erythematosus"

**Review and discussion for UVBACGs beside IFN-I stimulated genes (ISGs)**

**JUN**

Protein c-Jun binds with other proteins, such as c-Fos, to form an active transcription factor Activator protein 1 (AP-1) that participates in regulating gene expression in response to a variety of stimuli, e.g., cytokines, growth factors, stress, and bacterial/viral infections [1]. AP-1 controls a number of cellular processes including differentiation, proliferation, and apoptosis, both in physiological and pathologic status (inflammation, neoplasm, etc.) [1, 2].

c-JUN is involved in the autoimmune pathology of LE. A polymorphism in c-Jun has been linked to the SLE susceptibility [3]. The activation of c-Jun N-terminal kinases (JNK), coupled with a diminished ERK/JNK ratio in PBMCs, correlates with the severity of long-term organ damage in SLE patients [4]. The inflammation-triggered activation of JNK and p38 MAPK pathways in T and B lymphocytes could be the key mechanisms contributing to lymphocyte hyperactivity in SLE [5]. Decreased T cell DNA methyltransferase is observed in lupus, leading to higher c-Jun mRNA levels [6]. Additionally, SLE T cells showed elevated baseline c-Jun levels and phosphorylated upon activation [7]. Even the stimulation of IFN-inducible, lupus-susceptibility gene Ifi202 is partly via the JNK/c-Jun pathway in splenic B and T cells [8].

c-JUN can be activated by UV, manifesting in increased mRNA and protein expression, along with N-terminal phosphorylation. Exposure of human skin in vivo to low doses of UV activates EGFR and p21Ras, as well as further stimulates MAPK, ERK, JNK, and p38, which result in the induction of c-Jun in both fibroblasts and keratinocytes. The skin upregulation of c-Jun and functional activation of AP-1, promote the transcription of matrix metalloproteinases, while inhibit procollagen transcription and synthesis [9], resulting in diverse response including photoaging [9-12]. Katayama et al. also confirmed that UV activation induces c-JUN mRNA upregulation, potentiating SLE-associated inflammation [13].

Additionally, AP-1 activated by UV induces CYR61/CCN1 [14], Smad7 [15] in fibroblasts, and Snail in keratinocytes [16].

Our data reveal that JUN shows the strongest and most consistent correlation with SLEDAI among all genes, along with high expression levels of JUN-targeted genes MMPs. We found elevated MMP8 expression in LDGs, correlating with studies from Skopelja-Gardner et, al. [17, 18], suggesting that neutrophils may produce MMP8 extensively via AP-1, contributing to renal damage. Although JUN's differential expression between lupus and normal tissues is not as pronounced as that of some other candidate genes, it is strongly supported as a highly relevant gene for disease activity, and targeting it could effectively help prevent LE flare-ups. For instance, tRA can inhibit UV induction JUN and the AP-1 through its breakdown by the ubiquitin-proteasome pathway, thereby preventing photoaging [19]. This raises hope for incorporating such drugs into topical application for preventive measures in LE patients.

***PTTG1***

**PTTG1** is a regulatory protein in multiple biological processes such as chromosome stability, p53/TP53 pathway, and DNA repair. It is initially discovered and named as an oncogene because it significantly promotes tumorigenesis, angiogenesis and cancer progression, including B cell lymphoma [20]. To note, PTTG1 mRNA expression is critically enhanced by inflammatory cytokines during T cell proliferation and activation [21]. PTTG1 also performs an important anti-apoptotic role in UV-induced DNA damage and apoptosis and this role was mediated by JNK pathway [22]. In addition, PTTG1 risk loci are shared by several autoimmune diseases including SLE, particularly more active SLE [23-26]. In the clinical data from Katayama et al., PTTG1 mRNA expression was significantly increased after UVB photoprovocation in the skin of healthy, DLE, and SCLE volunteers [13].

Among our datasets, compared with normal controls, PTTG1 is highly expressed in LE samples, especially in samples of blood, CD4+ T, and B cells. In the skin PBMCs, monocytes and neutrophils, PTTG1 expression is also elevated. PTTG1 expression is highly correlated with SLE disease activity. Therefore, PTTG1 is a potential candidate for SLE activity evaluation and therapeutic target, as it plays important roles in promoting local and systemic autoimmunocyte activation and proliferation, as well as protecting epidermal cells against UV-induced DNA damage-mediated apoptosis.

***CAV1***

CAV1 encode a scaffolding protein Caveolin 1, which is the main component of the caveolae plasma membranes found in most cell types [27]. CAV1 has multiple functions such as tumorigenesis, insulin signaling, and tissue fibrosis, etc. among a number of diseases, i.e., cancer, pulmonary disease, neurodegeneration, muscular dystrophy, cardiomyopathy, atherosclerosis, and others [28].

Caveolae are types of rafts that are rich in proteins of the caveolin family (caveolin-1, -2 and -3) which present a distinct signaling platform. The dysfunction of lipid raft signaling involved in the pathogenesis of a variety of conditions, SLE included [29].

In most cases, caveolin family proteins are located on the plasma membrane and in the cytoplasm, the serum level of caveolin may not so specific to explain the exact cellular functions but can still present the association with SLE and activity [28]. It is found that anti-DNA antibodies enter cells through endocytosis of different caveolin proteins, so that this can partly explains that intranuclear antibodies are relatively common within lesions in lupus patients. Through their transfer via caveolin proteins and appearance and in both inflammatory and normal (non-inflamed) tissues, it can promote the spread of autoimmune responses [28, 30]. As such, CAV1 is also associated with the cell internalizing and nuclear localizing anti-DNA antibody, which can induce glomerular hypercellularity and proteinuria thus worsening the activity and severity of lupus [31].

Additionally, Caveolin-1 links integrin alpha subunits to the tyrosine kinase Fyn, which activates Shc and Grb2, thereby coupling integrins to the Ras-ERK pathway and promoting cell cycle [32]. The activation of ERK signaling is highlighted in LE diseases, which is also showed in our results.

In SLE, their dysfunction may promote the alteration of lipid rafts, thereby causing the aberrant activation of T and B cells [28, 33, 34]. The upregulation of CAV1 has been reported in three independent studies both in mRNA level in B cells [35, 36] and serum protein level [28, 35, 36]. CAV1 is also reported involved in the costimulatory signal essential for T-cell receptor (TCR)-mediated T-cell activation. In addition, its binding to DPP4 induces T-cell proliferation and NF-kappa-B activation in a T-cell receptor/CD3-dependent manner [37]. It has also been reported that CAV1 play a crucial role in immune cells for regulation of the cell signaling and inflammatory response [38], such as the Jak-Stat signaling pathway [39], as well as the activation of the PI3 kinase pathway [40].

UV radiation is an inducer of activation of CAV1 [41]. Although in our skin sample datasets (CLE), CAV1 did not appear significant changed, it is reported that UV irradiation inducing caveolin-1 phosphorylation thereby promoting in the activation of downstream pathways, MAPKs (P38, JNK), AKT, and AP-1 [42]. UV radiation can also upregulate the protein expression and phosphorylation of CAV1 in fibroblasts [43, 44], which in associated with stress-induced apoptosis. Caveolin-1 can led to inhibition of cisplatin and UVR-induced apoptosis in SCLC cells; and also, could decrease caspase-3 activity and increase the stability of Bcl-2 at the protein level [45].

In our results, CAV1 is highly upregulated in the samples of blood, PBMC, especially in T and B cells, and its expression in blood cell is correlated with SLE activity. Immunocyte enrichment estimation analysis shows that CAV1 expression is correlated with T and B cells, as well as epidermal keratinocyte, fibroblasts, which are consistence with the clues we found in the literature.

***HLA-F***

HLA-F is a non-classical major histocompatibility class Ib molecule that plays a role in immune surveillance, immune tolerance and inflammation, which might be involved in various physiological and pathological processes, such as viral infection, transplantation, pregnancy, cancer, and autoimmune diseases [46]. It has been reported HLA-F has a role in triggering or inhibiting NK cell functions through interacting with NK activating and inhibitory receptors, respectively [46-48]. It can also regulate the function of effector and CD8+ T cells [48, 49]. HLA-F is primarily localized in the cytoplasm but is also found on the surface of activated T, B, and NK cells. Its enhanced expression can lead to shedding into circulation as soluble HLA-F. HLA-F serves as a marker for activated lymphocytes and monocytes and its elevated levels in circulation can induce the production of anti-HLA-F autoantibodies. This process suggests that HLA-F and its soluble form could be novel immunomarkers for monitoring immune cell hyperactivity and disease activity in SLE [50]. HLA-F is expressed not only in immune cells but also in various tissues, including the placenta, tonsils, spleen, bladder, skin, thymus, and liver [50].

In our data, HLA-F is mostly highly elevated in LE skin tissues. Additionally, it is also increased in SLE blood, PBMCs, CD4, CD8, and monocytes. HLA-F is upregulated via UVB radiation [17]. The mRNA expression of HLA-F in blood immunocytes is correlated with SLE activity. The above information suggests that under LE inflammation, HLA-F can be induced by UV exposure. Immune cells may enhance HLA-F expression, and through interactions between HLA-F and its receptors during the disease process, it can contribute to the promotion of disease activity.

***HOPX***

HOPX, a homeodomain protein and transcription cofactor that regulate cell differentiation and function, is constitutively expressed in multiple immunocytes, such as CD4+, CD8+ T cells and B cells, as well as NK, NKT, and myeloid cells [51, 52]. It is associated with T/B cell development and linked to immune-related diseases [53-56]. It has been found in mouse that Hopx, induced in Tregs by DC, plays a role in maintaining Treg survival and functions by blocking intrinsic IL-2 production, and thereby mediate T cell unresponsiveness. This mechanism is crucial to sustaining immunotolerance in autoimmune disease [57]. In addition, Hopx can downregulate

the expression of the AP-1 complex (JUN/FOS), thereby inhibit T cell proliferation mediate T cell unresponsiveness to rechallenge with antigen [51].
In our study, we found HOPX is downregulated in blood and skin tissues, as well as multiple immunocytes in LE patients. HOPX is downexpressed in the skin (normal, DLE and SCLE) after UV exposure [13, 17]. Immunocyte enrichment estimation analysis showed that it can be expressed in multiple cellular components (including Treg cells) in both skin and blood. In addition, it is negatively correlated with SLE activity, which implies that it may be involved in immunotolerance and immunoinhibition. Notably, JUN is a positively-correlated UVBACG found in our study,
Since its aberrant-expression related mechanism has neither been found in LE nor in UV stimulation, there is necessity of further exploration. Notably, JUN is a UVBACG that shows a positive correlation in our study. Notably, JUN is a positively correlated UVBACG in our study, and previous research indicates that Hopx can downregulate JUN expression, thereby inhibiting T cell immunity [51]. This underscores the important role of interactions and balance between UVBACGs in the immunopathogenesis of LE.

***RPL3***

RPL3 is a component of the large subunit of cytoplasmic ribosomes, which is associated with virus infection [58], inherited hematopoietic abnormalities [59], and carcinogenesis [60], which can be linked to the IFN-1 immune response. In addition, RPL3 genetic alteration is associated with primary biliary cholangitis, an autoimmune liver disease with a strong hereditary component [61]. Moreover, the downregulation of RPL3 is associated with apoptosis activation in autoimmune disease benign multiple sclerosis [62].
RPL3 is significantly downregulated both in LE (DLE and SCLE) and normal skin tissues after UV radiation [13, 17]. In our datasets, RPL3 is also low-expressed in blood and skin, as well as multiple immunocytes. Its expression is negatively correlated with SLE activity. Therefore, this gene may be linked to apoptosis activation, as well as the

immunoresponses under exposure to UV. Since its aberrant-expression related mechanism has not been found in LE, there is necessity of further exploration.

**insights into our views on how to target UVBACGs for treating LE**

Except for novel pharmaceutical and therapeutic strategy mining, daily protection from UVB radiation is crucial. The guidelines for treating SLE strongly advise educating patients on using broad-spectrum sunscreens for effective protection against UV exposure [63]. However, clinical studies review a significant lack of knowledge among patients and parents of SLE children, as well as rheumatologists regarding sunscreen recommendations [64-66].

Additionally, clinical guidance on the use of indoor lamps remains uncertain. A number of light bulbs found in homes and workplaces (e.g., manicures) emit low doses of UV radiation. Although the intensity is much lower than that of sunlight, the extended exposure time lasting for hours and repeated daily use could potentially lead to a notable accumulation of damage [67, 68].

Thus, avoidance of daily UV radiation is arduous and, admittedly, there are so many advantages of natural sunlight to human body. For instance, low-dose UVA1 irradiation can be used to treat LE and can significantly decrease the disease activity [67, 69, 70]. Additionally, vitamin D is crucial for patients with LE. Supplementing with vitamin D has proven effective in reducing disease activity, as it plays a key role in immunoregulation [71-74]. Unfortunately, individuals with LE, especially those with SLE, commonly face a deficiency in vitamin D, primarily sourced from UVB exposure [71, 72, 74].

Hence, we tentatively propose holistic therapeutic approaches that include: 1) systemic administration of regular and/or UVBACG-targeted medications; 2) the use of sunscreen along with vitamin D supplements; and 3) topical applications of UVBACG-targeted agents. Regarding the first point, targeted medications should gradually replace regular ones to minimize severe side effects. Emphasizing the third point is crucial, given that the skin is the “primary crime scene” of LE. However, till now, there are only a few clinical trials or case reports available. Topical LE treatment starts with corticosteroid application, but prolonged use leads to unavoidable, severe side effects [75, 76]. Calcineurin inhibitors, primarily tacrolimus, were tested in clinical observations and trials. While 50% of patients responded, DLE and SCLE lesions showed lower responsiveness [75, 77, 78]. Recent three years, topical JAK inhibitors, crucial in restraining JAK-STAT pathways involved in the autocrine loop for IFN-I, have been gradually incorporated into clinical observations. Examples include Delgocitinib for an SCLE patient [76], JAK1/2 inhibitor ruxolitinib for an SLE/DLE patient with scalp lesions [79], and JAK1/3 inhibitor tofacitinib for a periorbital DLE patient [80]. However, some trials did not yield significant improvement, e.g., R333 (JAK/SYK inhibitor) for DLE, [81], GSK2646264 (SYK inhibitor) for active CLE [82]. Additional adjuvant medications, such as niacinamide for DLE [83], clindamycin for CCLE [84], R-salbutamol for DLE [85], and halobetasol propionate for DLE[77], have shown some benefits. Our study suggests exploring more topical medications targeting UVBACGs, and related cells/pathways. Due to limited experience and evidence, further large cohort studies are necessary to identify more effective, safe, and tolerable treatments.

Finally, provided that the presence or absence of microbes determines whether UVB induces a pro-inflammatory environment or an immunosuppressive state in the skin. Thus, modulating the skin microbiota to block UVB-induced autoimmune activation is another potential therapeutic strategy [86].

**This study has some limitations:**

1) Confirming whether skin lesions from sunlight exposure or UV radiation are synonymous with UVB phototesting-defined photosensitivity requires more thorough examination, since UVB is only a part of sunlight. In the current study, we employed a substantial number of non-artificially irradiated LE samples, thereby partially addressing this limitation. The effects of other sunlight or UV wavelengths on LE require further investigation.

2) We did not account for a more extended observation period, since short-term UVB-MED may not comprehensively represent UVB photosensitivity. Sanders et al. suggest that trials with observation periods of up to 8 weeks reveal a significantly higher

number of photosensitive patients, thus recommending the implementation of phototesting with an extended protocol and prolonged observation periods [87].

3) More extensive clinical studies are required to validate the correlation between UVB-MED (and risk-evaluation models) and SLE activity. Additionally, determining the clinical significance reference value, possibly around 56.525 mJ/cm2, needs further investigation.

4) To comprehensively gauge the severity and activity of SLE, utilizing various evaluation systems (e.g., PGA, BILAG, ECLAM, SLAM) alongside SLEDAI is crucial, considering the diverse advantages and drawbacks of each.

5) This study did not collect clinical data and UVB-MED from CLE patients, thus hindering the analysis of CLE photosensitivity. Further research will focus on the correlation between the RCLASI score of CLE patients and UVB sensitivity.

6) Heterogeneity in transcriptomic datasets due to factors like ethnicity, lifestyles, and therapeutic strategies does exists. Thus, we select genes consistently expressed across most datasets.

7） The samples in this study were not collected from the same individual at different stages. Our study focuses on association and correlation, providing strong clues to the UVB-induced LE pathogenesis. However, it cannot establish a causal relationship between UV provocation, autoimmune activation, and LE activity. Similarly, the specific sequence and causal relationship of UVBACGs, UVBACG/SLEDAI-correlated genes, and related pathways' activation cannot be definitively elucidated.

8) Our skin samples are limited to CLE, while blood and immunocytes are obtained from SLE. As a result, we are unable to compare CLE and SLE within the same patient and cannot make assumptions about the progression from CLE to SLE.

9) Limitations exist in both functional enrichment and the estimation of cell enrichment/abundance (xCell, CIBERSORT, and ImmuCellAI), relying on further analysis of single-cell sequencing or histological studies.

**References**

1. Wisdom, R., R.S. Johnson, and C. Moore, *c-Jun regulates cell cycle progression*

*and apoptosis by distinct mechanisms.* EMBO J, 1999. **18**(1): p. 188-97.
2. Shaulian, E. and M. Karin, *AP-1 in cell proliferation and survival.* Oncogene, 2001. **20**(19): p. 2390-400.
3. Ma, Y., et al., *Association of c-Jun gene polymorphism with susceptibility to systemic lupus erythematosus in a Chinese population.* DNA Cell Biol, 2012. **31**(7): p. 1274-8.
4. Bloch, O., et al., *Increased ERK and JNK activation and decreased ERK/JNK ratio are associated with long-term organ damage in patients with systemic lupus erythematosus.* Rheumatology (Oxford), 2014. **53**(6): p. 1034-42.
5. Wong, C.K., et al., *Activation profile of intracellular mitogen-activated protein kinases in peripheral lymphocytes of patients with systemic lupus erythematosus.* J Clin Immunol, 2009. **29**(6): p. 738-46.
6. Yang, J., et al., *Effect of mitogenic stimulation and DNA methylation on human T cell DNA methyltransferase expression and activity.* J Immunol, 1997. **159**(3): p. 1303-9.
7. Ghosh, D., G.C. Tsokos, and V.C. Kyttaris, *c-Jun and Ets2 proteins regulate expression of spleen tyrosine kinase in T cells.* J Biol Chem, 2012. **287**(15): p. 11833-41.
8. Chen, J., R. Panchanathan, and D. Choubey, *Stimulation of T cells up-regulates expression of Ifi202, an interferon-inducible lupus susceptibility gene, through activation of JNK/c-Jun pathway.* Immunol Lett, 2008. **118**(1): p. 13-20.
9. Fisher, G.J., et al., *c-Jun-dependent inhibition of cutaneous procollagen transcription following ultraviolet irradiation is reversed by all-trans retinoic acid.* J Clin Invest, 2000. **106**(5): p. 663-70.
10. Fisher, G.J., et al., *Retinoic acid inhibits induction of c-Jun protein by ultraviolet radiation that occurs subsequent to activation of mitogen-activated protein kinase pathways in human skin in vivo.* J Clin Invest, 1998. **101**(6): p. 1432-40.
11. Fisher, G.J., et al., *Mechanisms of photoaging and chronological skin aging.* Arch Dermatol, 2002. **138**(11): p. 1462-70.
12. Angel, P., A. Szabowski, and M. Schorpp-Kistner, *Function and regulation of AP-1 subunits in skin physiology and pathology.* Oncogene, 2001. **20**(19): p. 2413-23.
13. Katayama, S., et al., *Delineating the Healthy Human Skin UV Response and Early Induction of Interferon Pathway in Cutaneous Lupus Erythematosus.* J Invest Dermatol, 2019. **139**(9): p. 2058-2061 e4.
14. Quan, T., et al., *Ultraviolet irradiation induces CYR61/CCN1, a mediator of collagen homeostasis, through activation of transcription factor AP-1 in human skin fibroblasts.* J Invest Dermatol, 2010. **130**(6): p. 1697-706.
15. Quan, T., et al., *Ultraviolet irradiation induces Smad7 via induction of transcription factor AP-1 in human skin fibroblasts.* J Biol Chem, 2005. **280**(9): p. 8079-85.
16. Li, Y., et al., *UV irradiation induces Snail expression by AP-1 dependent mechanism in human skin keratinocytes.* J Dermatol Sci, 2010. **60**(2): p. 105-13.
17. Skopelja-Gardner, S., et al., *The early local and systemic Type I interferon*

*responses to ultraviolet B light exposure are cGAS dependent.* Sci Rep, 2020. **10**(1): p. 7908.

18. Skopelja-Gardner, S., et al., *Acute skin exposure to ultraviolet light triggers neutrophil-mediated kidney inflammation.* Proc Natl Acad Sci U S A, 2021. **118**(3).
19. Fisher, G.J., et al., *Molecular mechanisms of photoaging in human skin in vivo and their prevention by all-trans retinoic acid.* Photochem Photobiol, 1999. **69**(2): p. 154-7.
20. Chiriva-Internati, M., et al., *The pituitary tumor transforming gene 1 (PTTG-1): an immunological target for multiple myeloma.* J Transl Med, 2008. **6**: p. 15.
21. Stoika, R. and S. Melmed, *Expression and function of pituitary tumour transforming gene for T-lymphocyte activation.* Br J Haematol, 2002. **119**(4): p. 1070-4.
22. Lai, Y., et al., *The important anti-apoptotic role and its regulation mechanism of PTTG1 in UV-induced apoptosis.* J Biochem Mol Biol, 2007. **40**(6): p. 966-72.
23. Chung, S.A., et al., *Differential genetic associations for systemic lupus erythematosus based on anti-dsDNA autoantibody production.* PLoS Genet, 2011. **7**(3): p. e1001323.
24. Gateva, V., et al., *A large-scale replication study identifies TNIP1, PRDM1, JAZF1, UHRF1BP1 and IL10 as risk loci for systemic lupus erythematosus.* Nat Genet, 2009. **41**(11): p. 1228-33.
25. Fang, K., K. Zhang, and J. Wang, *Network-assisted analysis of primary Sjogren's syndrome GWAS data in Han Chinese.* Sci Rep, 2015. **5**: p. 18855.
26. Mathieu, A., et al., *Genetics of psoriasis and psoriatic arthritis.* Reumatismo, 2007. **59 Suppl 1**: p. 25-7.
27. Vargas, L., et al., *Functional interaction of caveolin-1 with Bruton's tyrosine kinase and Bmx.* J Biol Chem, 2002. **277**(11): p. 9351-7.
28. Li, M., et al., *Expression of caveolin family proteins in serum of patients with systemic lupus erythematosus.* Lupus, 2021. **30**(11): p. 1819-1828.
29. Michel, V. and M. Bakovic, *Lipid rafts in health and disease.* Biol Cell, 2007. **99**(3): p. 129-40.
30. Golan, T.D., et al., *The penetrating potential of autoantibodies into live cells in vitro coincides with the in vivo staining of epidermal nuclei.* Lupus, 1997. **6**(1): p. 18-26.
31. Yanase, K. and M.P. Madaio, *Nuclear localizing anti-DNA antibodies enter cells via caveoli and modulate expression of caveolin and p53.* J Autoimmun, 2005. **24**(2): p. 145-51.
32. Wary, K.K., et al., *A requirement for caveolin-1 and associated kinase Fyn in integrin signaling and anchorage-dependent cell growth.* Cell, 1998. **94**(5): p. 625-34.
33. Deng, G.M. and G.C. Tsokos, *Cholera toxin B accelerates disease progression in lupus-prone mice by promoting lipid raft aggregation.* J Immunol, 2008. **181**(6): p. 4019-26.
34. Vasquez, A., et al., *Altered recruitment of Lyn, Syk and ZAP-70 into lipid rafts of*

*activated B cells in Systemic Lupus Erythematosus.* Cell Signal, 2019. **58**: p. 9-19.
35. Udhaya Kumar, S., et al., *Dysregulation of Signaling Pathways Due to Differentially Expressed Genes From the B-Cell Transcriptomes of Systemic Lupus Erythematosus Patients - A Bioinformatics Approach.* Front Bioeng Biotechnol, 2020. **8**: p. 276.
36. Fan, H., et al., *Gender differences of B cell signature in healthy subjects underlie disparities in incidence and course of SLE related to estrogen.* J Immunol Res, 2014. **2014**: p. 814598.
37. Ohnuma, K., et al., *Caveolin-1 triggers T-cell activation via CD26 in association with CARMA1.* J Biol Chem, 2007. **282**(13): p. 10117-10131.
38. Aravamudan, B., et al., *Caveolin-1 knockout mice exhibit airway hyperreactivity.* Am J Physiol Lung Cell Mol Physiol, 2012. **303**(8): p. L669-81.
39. Guo, C.J., et al., *Involvement of caveolin-1 in the Jak-Stat signaling pathway and infectious spleen and kidney necrosis virus infection in mandarin fish (Siniperca chuatsi).* Mol Immunol, 2011. **48**(8): p. 992-1000.
40. Sud, N., D.A. Wiseman, and S.M. Black, *Caveolin 1 is required for the activation of endothelial nitric oxide synthase in response to 17beta-estradiol.* Mol Endocrinol, 2010. **24**(8): p. 1637-49.
41. Wang, Z., et al., *Caveolin-1, a stress-related oncotarget, in drug resistance.* Oncotarget, 2015. **6**(35): p. 37135-50.
42. Wang, Q., et al., *Extracellular matrix activity and caveolae events contribute to cell surface receptor activation that leads to MAP kinase activation in response to UV irradiation in cultured human keratinocytes.* Int J Mol Med, 2005. **15**(4): p. 633-40.
43. Volonte, D., et al., *Expression of caveolin-1 induces premature cellular senescence in primary cultures of murine fibroblasts.* Mol Biol Cell, 2002. **13**(7): p. 2502-17.
44. Volonte, D., et al., *Cellular stress induces the tyrosine phosphorylation of caveolin-1 (Tyr(14)) via activation of p38 mitogen-activated protein kinase and c-Src kinase. Evidence for caveolae, the actin cytoskeleton, and focal adhesions as mechanical sensors of osmotic stress.* J Biol Chem, 2001. **276**(11): p. 8094-103.
45. Yang, X., et al., *Higher expression of Caveolin-1 inhibits human small cell lung cancer (SCLC) apoptosis in vitro.* Cancer Invest, 2012. **30**(6): p. 453-62.
46. Lin, A. and W.H. Yan, *The Emerging Roles of Human Leukocyte Antigen-F in Immune Modulation and Viral Infection.* Front Immunol, 2019. **10**: p. 964.
47. Garcia-Beltran, W.F., et al., *Open conformers of HLA-F are high-affinity ligands of the activating NK-cell receptor KIR3DS1.* Nat Immunol, 2016. **17**(9): p. 1067-74.
48. Goodridge, J.P., et al., *HLA-F and MHC class I open conformers are ligands for NK cell Ig-like receptors.* J Immunol, 2013. **191**(7): p. 3553-62.
49. Goodridge, J.P., et al., *HLA-F and MHC-I open conformers cooperate in a MHC-I antigen cross-presentation pathway.* J Immunol, 2013. **191**(4): p. 1567-77.

50. Jucaud, V., et al., *Serum antibodies to human leucocyte antigen (HLA)-E, HLA-F and HLA-G in patients with systemic lupus erythematosus (SLE) during disease flares: Clinical relevance of HLA-F autoantibodies.* Clin Exp Immunol, 2016. **183**(3): p. 326-40.
51. Hawiger, D., et al., *The transcription cofactor Hopx is required for regulatory T cell function in dendritic cell-mediated peripheral T cell unresponsiveness.* Nat Immunol, 2010. **11**(10): p. 962-8.
52. Bourque, J., et al., *Landscape of Hopx expression in cells of the immune system.* Heliyon, 2021. **7**(11): p. e08311.
53. Albrecht, I., et al., *Persistence of effector memory Th1 cells is regulated by Hopx.* Eur J Immunol, 2010. **40**(11): p. 2993-3006.
54. Opejin, A., et al., *A Two-Step Process of Effector Programming Governs CD4(+) T Cell Fate Determination Induced by Antigenic Activation in the Steady State.* Cell Rep, 2020. **33**(8): p. 108424.
55. Descatoire, M., et al., *Identification of a human splenic marginal zone B cell precursor with NOTCH2-dependent differentiation properties.* J Exp Med, 2014. **211**(5): p. 987-1000.
56. Kogan, V., et al., *Genetic-Epigenetic Interactions in Asthma Revealed by a Genome-Wide Gene-Centric Search.* Hum Hered, 2018. **83**(3): p. 130-152.
57. Jones, A., et al., *Peripherally Induced Tolerance Depends on Peripheral Regulatory T Cells That Require Hopx To Inhibit Intrinsic IL-2 Expression.* J Immunol, 2015. **195**(4): p. 1489-97.
58. Labruyere, E., et al., *Crosstalk between Entamoeba histolytica and the human intestinal tract during amoebiasis.* Parasitology, 2019. **146**(9): p. 1140-1149.
59. Choijilsuren, H.B., Y. Park, and M. Jung, *Mechanisms of somatic transformation in inherited bone marrow failure syndromes.* Hematology Am Soc Hematol Educ Program, 2021. **2021**(1): p. 390-398.
60. Zhang, X., et al., *DUOX2 promotes the progression of colorectal cancer cells by regulating the AKT pathway and interacting with RPL3.* Carcinogenesis, 2021. **42**(1): p. 105-117.
61. Qiu, F., et al., *A genome-wide association study identifies six novel risk loci for primary biliary cholangitis.* Nat Commun, 2017. **8**: p. 14828.
62. Achiron, A., et al., *Suppressed RNA-polymerase 1 pathway is associated with benign multiple sclerosis.* PLoS One, 2012. **7**(10): p. e46871.
63. Fanouriakis, A., et al., *2019 update of the EULAR recommendations for the management of systemic lupus erythematosus.* Ann Rheum Dis, 2019. **78**(6): p. 736-745.
64. Akamine, K.L., et al., *Trends in sunscreen recommendation among US physicians.* JAMA Dermatol, 2014. **150**(1): p. 51-5.
65. Pratt, A., et al., *Sunscreen knowledge amongst rheumatologists: Finding the gap.* Lupus, 2021. **30**(2): p. 360-362.
66. Janthongsri, T., W. Wisuthsarewong, and R. Nitiyarom, *Photoprotective habits in children with systemic lupus erythematosus.* Lupus, 2020. **29**(8): p. 964-969.
67. Klein, R.S., et al., *The risk of ultraviolet radiation exposure from indoor lamps*

*in lupus erythematosus.* Autoimmun Rev, 2009. **8**(4): p. 320-4.
68. Keyes, E., et al., *Ultraviolet light exposure from manicures in cutaneous lupus erythematosus.* Rheumatology (Oxford), 2022. **61**(2): p. e38-e39.
69. McGrath, H., Jr., *Ultraviolet-A1 irradiation therapy for systemic lupus erythematosus.* Lupus, 2017. **26**(12): p. 1239-1251.
70. Li, Q., et al., *An Update on the Pathogenesis of Skin Damage in Lupus.* Curr Rheumatol Rep, 2020. **22**(5): p. 16.
71. Schneider, L., et al., *Vitamin D and systemic lupus erythematosus: state of the art.* Clin Rheumatol, 2014. **33**(8): p. 1033-8.
72. Irfan, S.A., et al., *Effects of Vitamin D on Systemic Lupus Erythematosus Disease Activity and Autoimmunity: A Systematic Review and Meta-Analysis.* Cureus, 2022. **14**(6): p. e25896.
73. Cutillas-Marco, E., et al., *Vitamin D and cutaneous lupus erythematosus: effect of vitamin D replacement on disease severity.* Lupus, 2014. **23**(7): p. 615-23.
74. Cusack, C., et al., *Photoprotective behaviour and sunscreen use: impact on vitamin D levels in cutaneous lupus erythematosus.* Photodermatol Photoimmunol Photomed, 2008. **24**(5): p. 260-7.
75. Kanekura, T., et al., *Efficacy of topical tacrolimus for treating the malar rash of systemic lupus erythematosus.* Br J Dermatol, 2003. **148**(2): p. 353-6.
76. Maruyama, A. and N. Katoh, *Subacute cutaneous lupus erythematosus successfully treated with topical delgocitinib.* J Dermatol, 2023. **50**(3): p. e110-e111.
77. Barua, D.P., et al., *Comparison of effectiveness of topical tacrolimus 0.1% vs topical halobetasol propionate 0.05% as an add-on to oral hydroxychloroquine in discoid lupus erythematosus.* Dermatol Ther, 2021. **34**(1): p. e14675.
78. Lampropoulos, C.E. and D.P. D'Cruz, *Topical calcineurin inhibitors in systemic lupus erythematosus.* Ther Clin Risk Manag, 2010. **6**: p. 95-101.
79. Park, J.J., A.J. Little, and M.D. Vesely, *Treatment of cutaneous lupus with topical ruxolitinib cream.* JAAD Case Rep, 2022. **28**: p. 133-135.
80. Mazori, D.R., et al., *Use of Tofacitinib, 2%, Ointment for Periorbital Discoid Lupus Erythematosus.* JAMA Dermatol, 2021. **157**(7): p. 880-882.
81. Presto, J.K., et al., *Computerized planimetry to assess clinical responsiveness in a phase II randomized trial of topical R333 for discoid lupus erythematosus.* Br J Dermatol, 2018. **178**(6): p. 1308-1314.
82. Walker, A., et al., *Safety, pharmacokinetics and pharmacodynamics of a topical SYK inhibitor in cutaneous lupus erythematosus: A double-blind Phase Ib study.* Exp Dermatol, 2021. **30**(11): p. 1686-1692.
83. Nouh, A.H., et al., *Topical niacinamide (Nicotinamide) treatment for discoid lupus erythematosus (DLE): A prospective pilot study.* J Cosmet Dermatol, 2023. **22**(5): p. 1647-1657.
84. Newman, A.J., et al., *Chronic cutaneous lupus erythematosus and topical clindamycin.* BMJ Case Rep, 2018. **2018**.
85. Jemec, G.B., et al., *A randomized controlled trial of R-salbutamol for topical treatment of discoid lupus erythematosus.* Br J Dermatol, 2009. **161**(6): p.

1365-70.
86. Patra, V., et al., *Skin Microbiome Modulates the Effect of Ultraviolet Radiation on Cellular Response and Immune Function.* iScience, 2019. **15**: p. 211-222.
87. Sanders, C.J., et al., *Photosensitivity in patients with lupus erythematosus: a clinical and photobiological study of 100 patients using a prolonged phototest protocol.* Br J Dermatol, 2003. **149**(1): p. 131-7.
