## Supplementary Figures for "Exploring the Correlation Between UVB Sensitivity and SLE Activity: Insights into UVB-Driven Pathogenesis in Lupus Erythematosus"

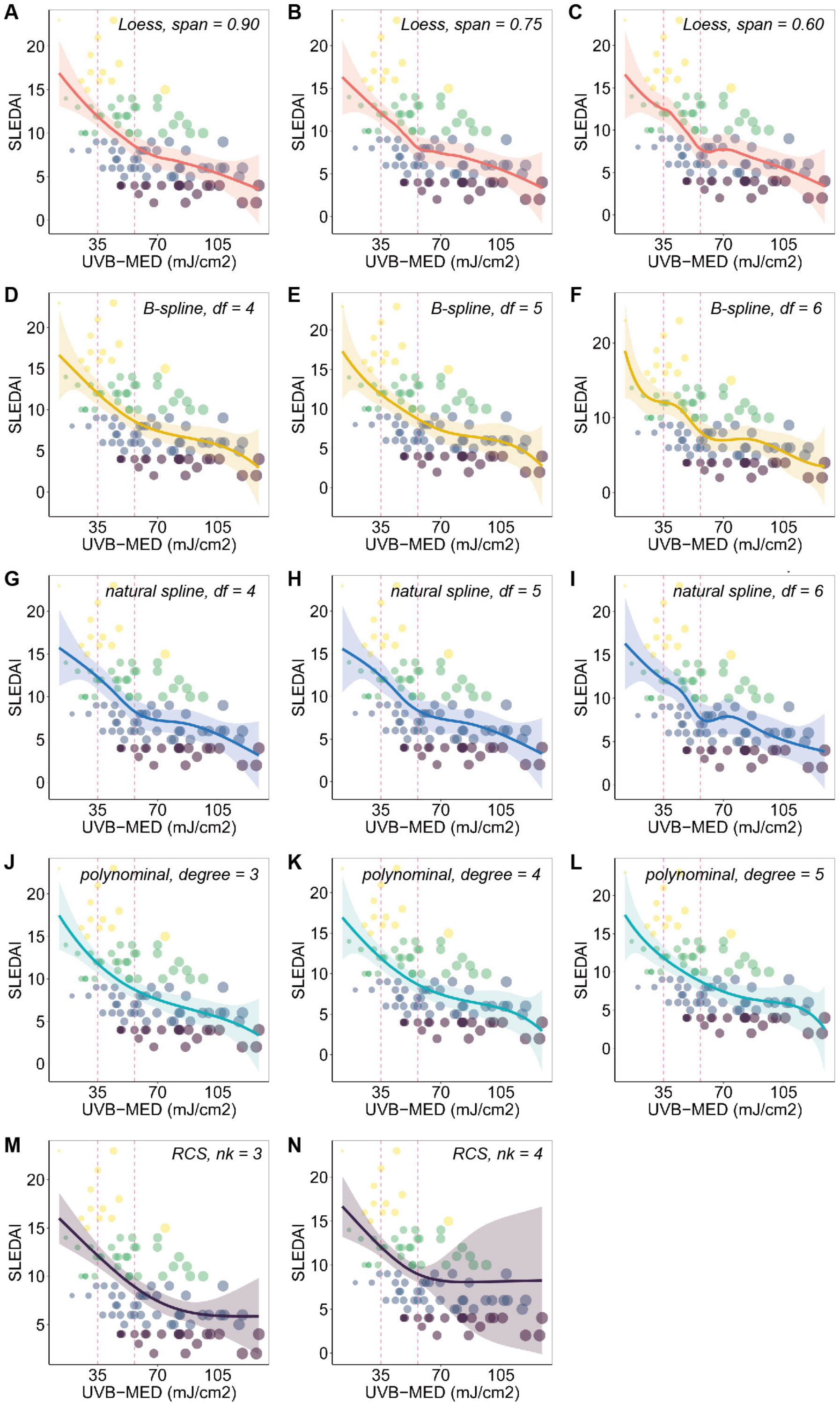


**Fig. S1. Non-linear regressions between SLEDAI and UVB-MED.** Illustration of non-linear regressions employing gradually varied parameters for each regression method: Loess (locally weighted regression), B-spline (basis polynomial spline), RCS (restricted cubic spline), and Polynomial regression. The dashed lines in all figures indicate the thresholds of 35 and 56.525 mJ/cm$^2$.

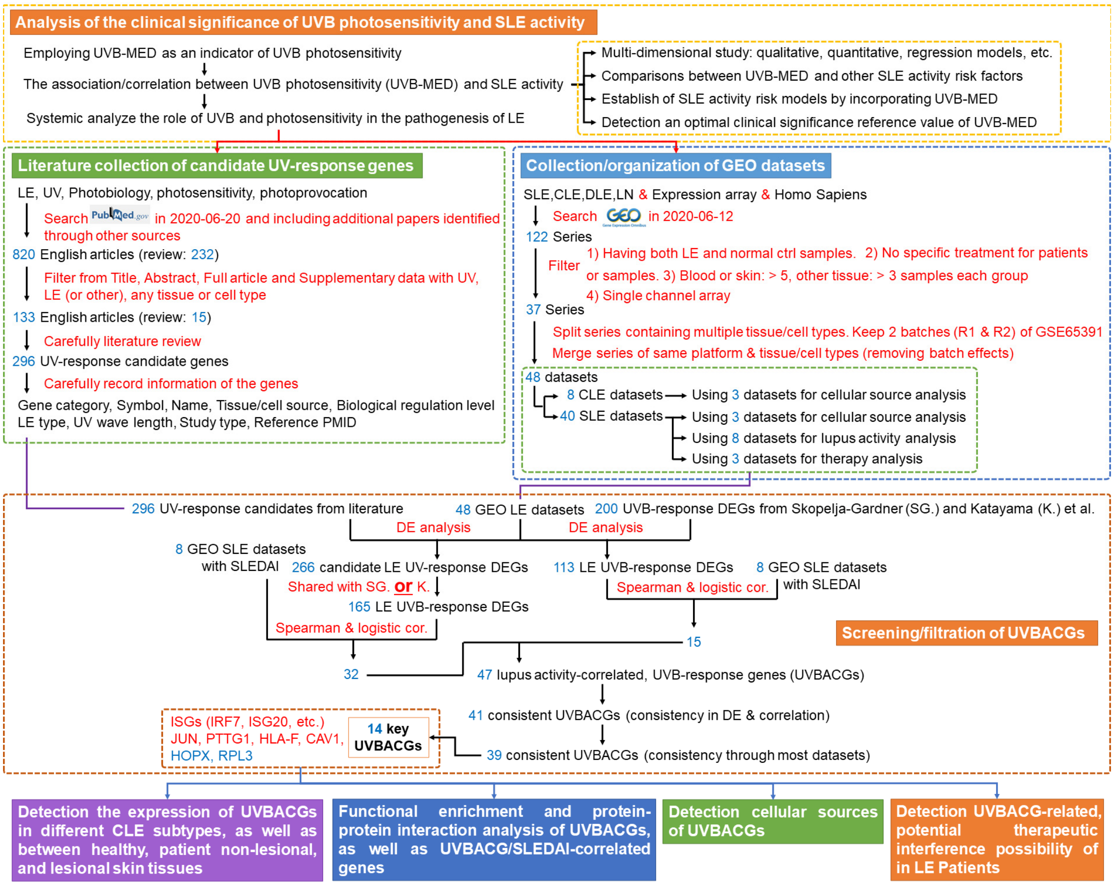


**Fig. S2. Flow chart summarizing the structure and content of the present study.** This comprehensive flow chart illustrates the overall structure and content of the study. Abbreviations: LE (lupus erythematosus), DE (differential expression), DEGs (differentially expressed genes).

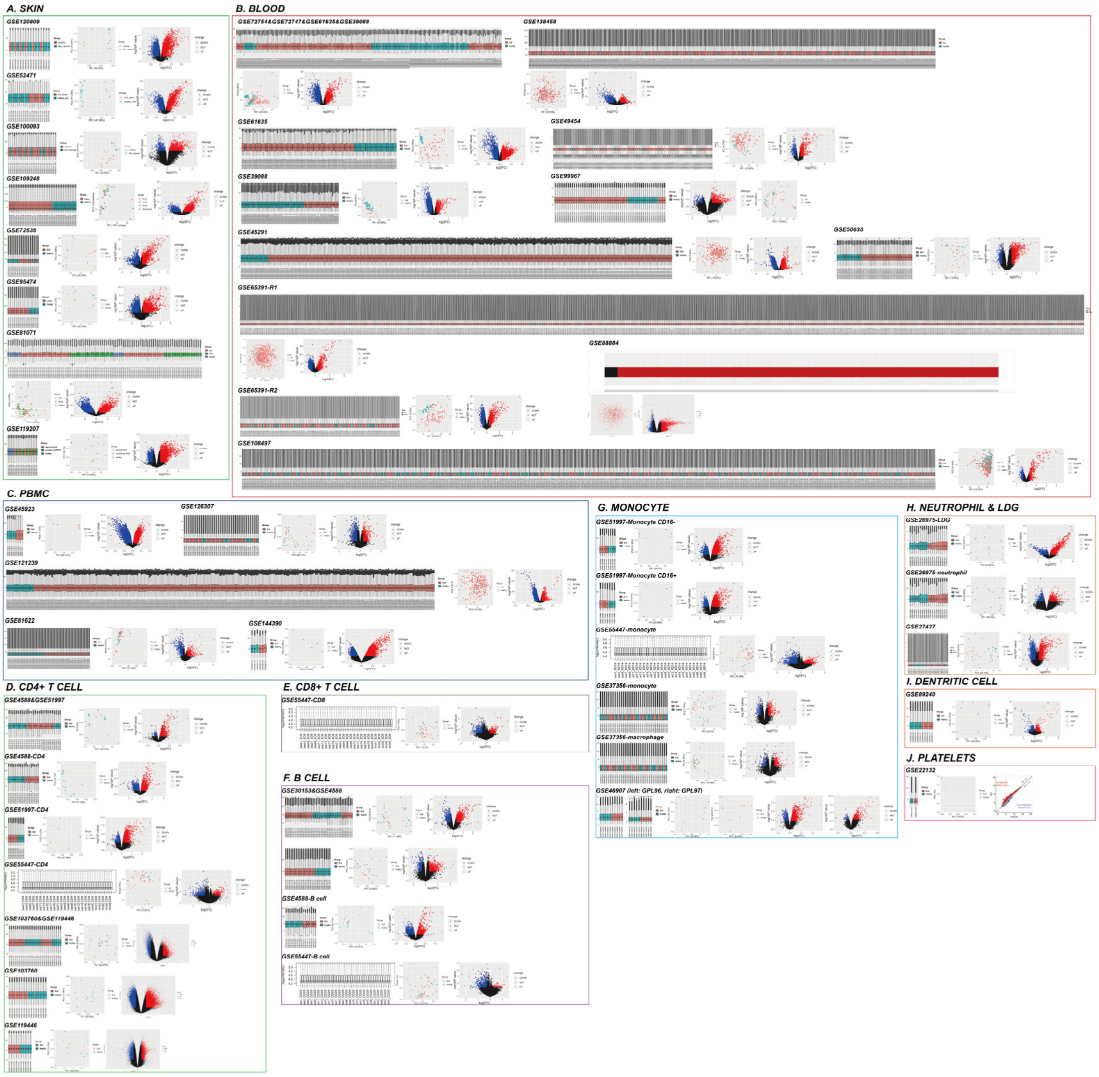


**Fig. S3**. Boxplots, PCA plots, and volcano plots illustrate the data distribution, group differences, and differentially expressed genes across the 48 GEO SLE datasets included in this study.

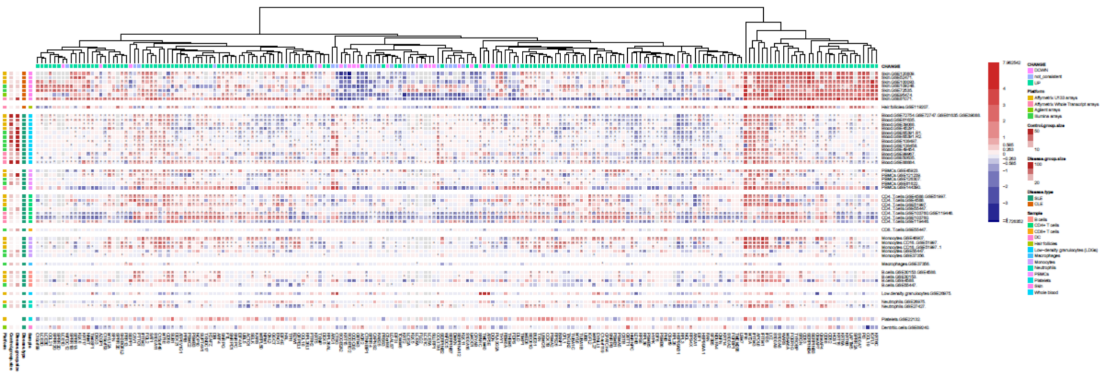

**Fig. S4**. A heatmap displaying differential gene expression (fold change) for LE samples compared to normal samples across the 200 UVB-response candidate DEGs identified by Skopelja-Gardner et al. and Katayama et al., in addition to the 296-gene list.

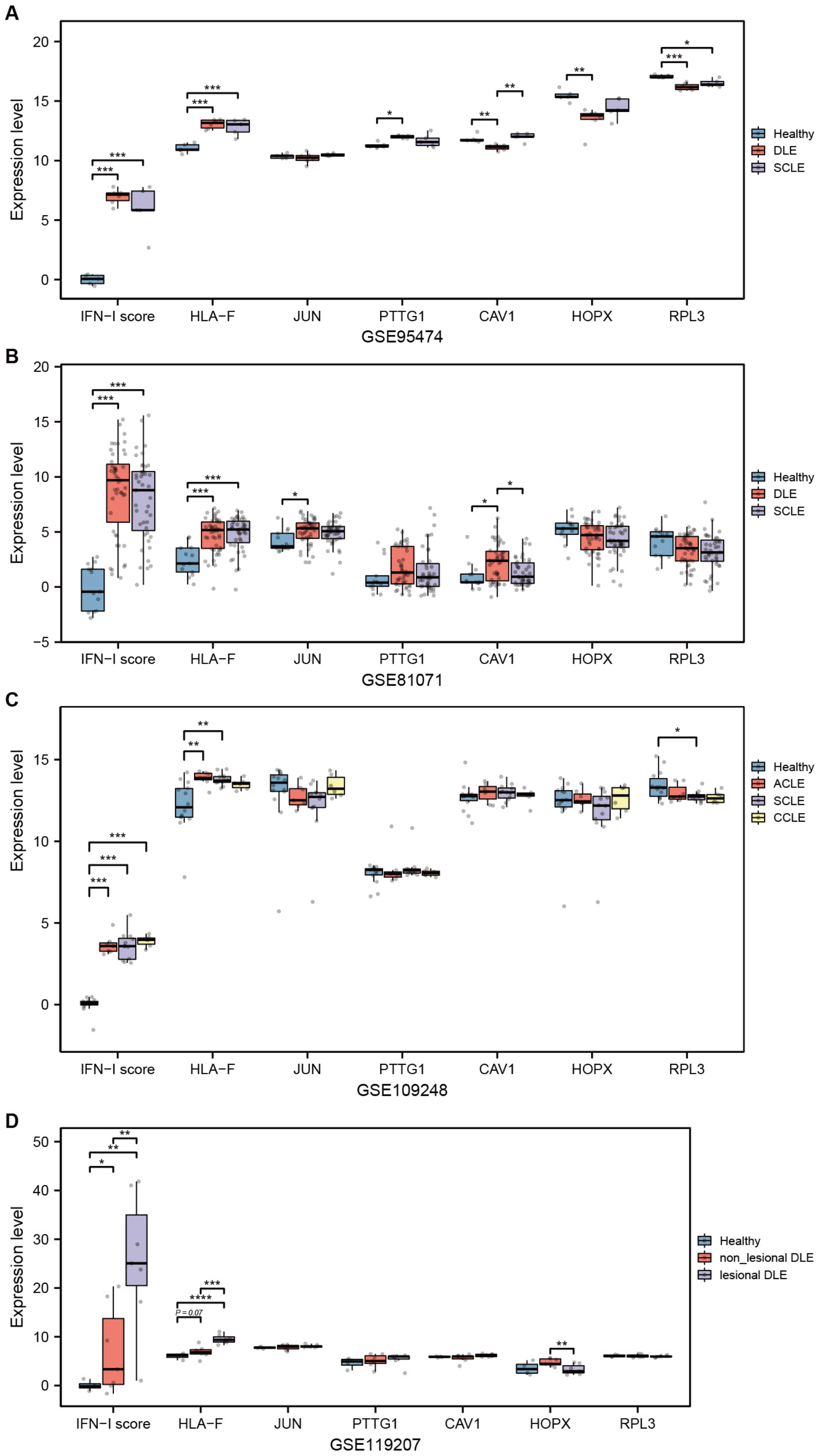


**Fig. S5**. Differential expression of UVBACGs in different CLE subtypes (**A-C**), as well as between healthy, patient non-lesional, and lesional skin follicle tissues (**D**). * P < 0.05, ** P < 0.01, *** P < 0.001, **** P < 0.0001.

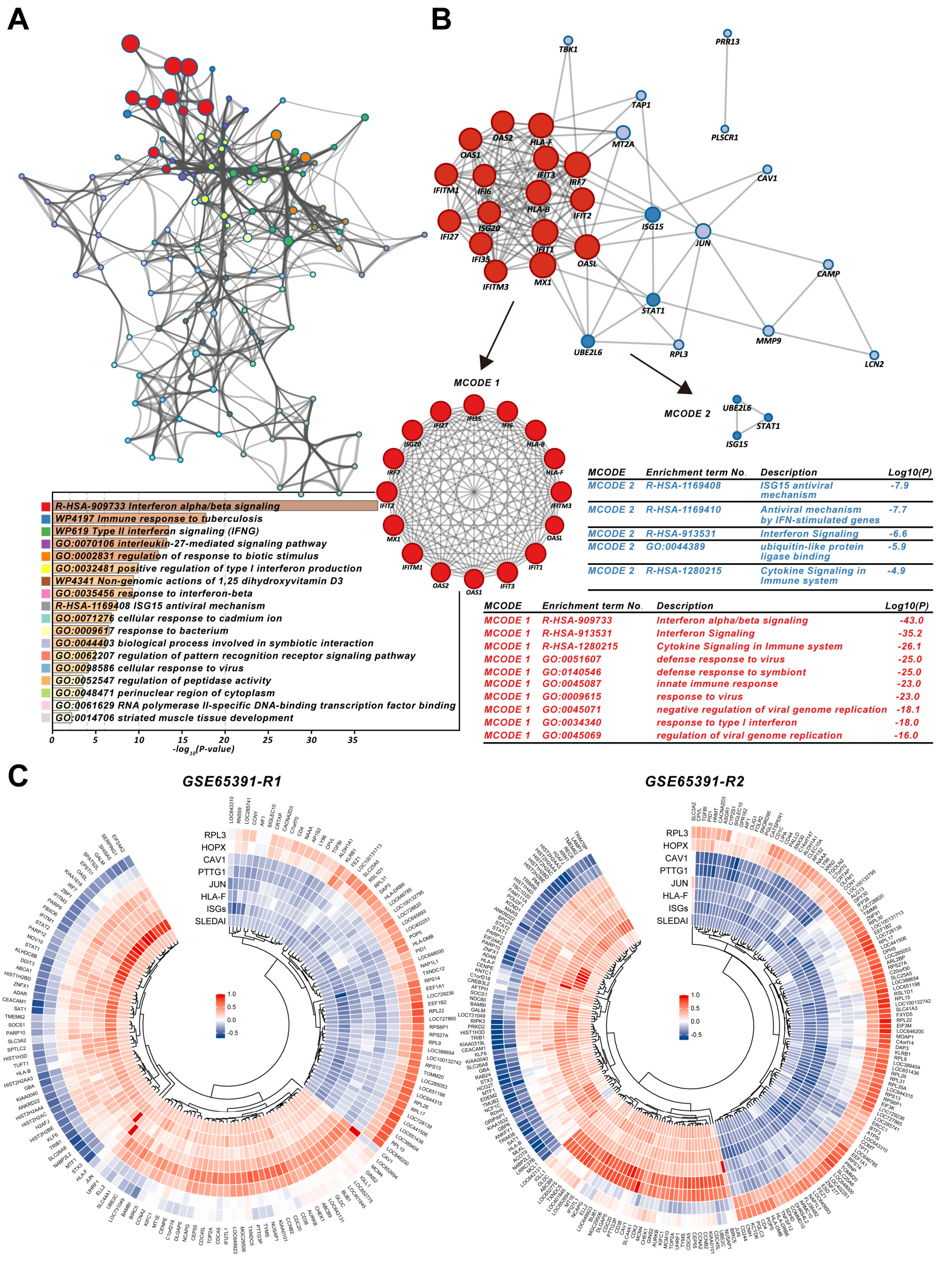


**Fig. S6**. **A, B**) Functional enrichment and protein-protein interaction analysis of 39 consistent UVBACGs showed significant features of interferon signaling (especially IFN

I), response to virus infection, and activation of type I immunity related intracellular pathways. **C**) Circular heat maps showing the expression correlation of key UVBACG/SLEDAI-correlated genes with key UVBACGs (Pearson) and SLEDAI (Spearman), using GSE65391-R1 and R2 datasets.

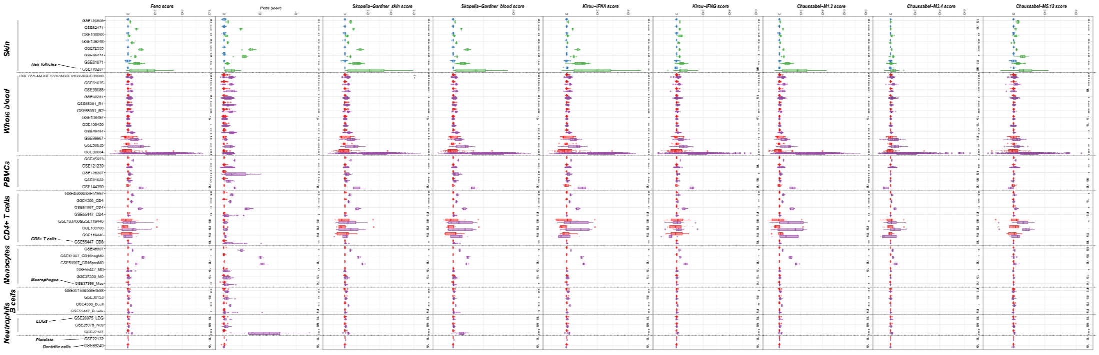


**Fig. S7**. Comparisons of IFN signature scores between LE samples (CLE in green and SLE in purple) and control samples across all datasets, based on published IFN signature genes [Feng score, Skopelja-Gardner skin and blood score, Kirou IFNA (IFNα) and IFNG (IFNγ) score, Chaussasel M1.2 (IFNα), M3.4 (IFNβ) and M5.12 (IFNγ) scores]. The results indicate that most tissue/cell samples, including skin, blood, PBMCs, T/B cells, monocytes/macrophages, and platelets, exhibit the highest differences in the IFNα signature, followed by IFNβ, and then IFNγ.

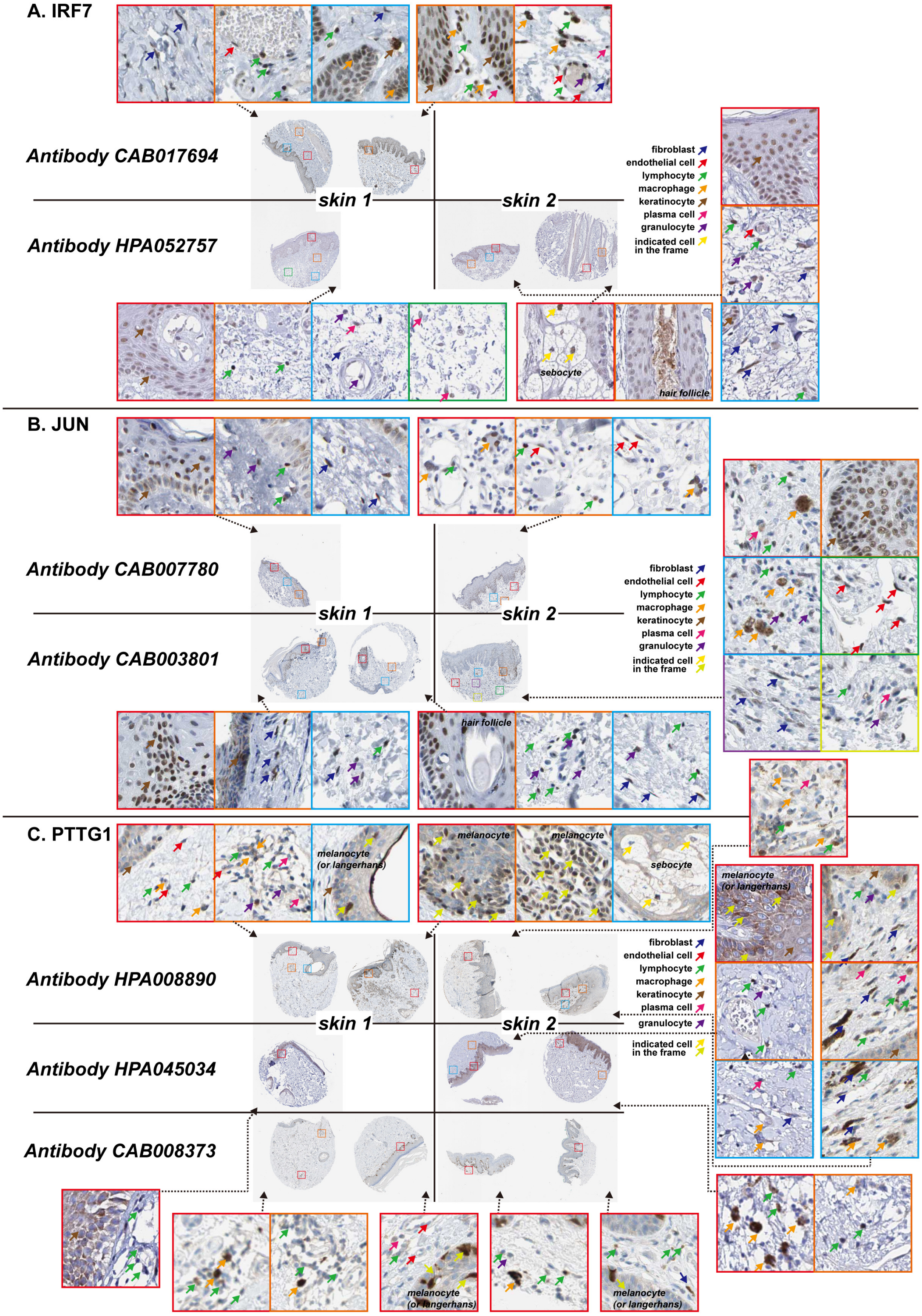


**Fig. S8**. Potential cellular source evidence of UVBACG expression by immunohistochemistry staining on skin tissue microslides from Human Protein Atlas (HPA) (https://www.proteinatlas.org/).

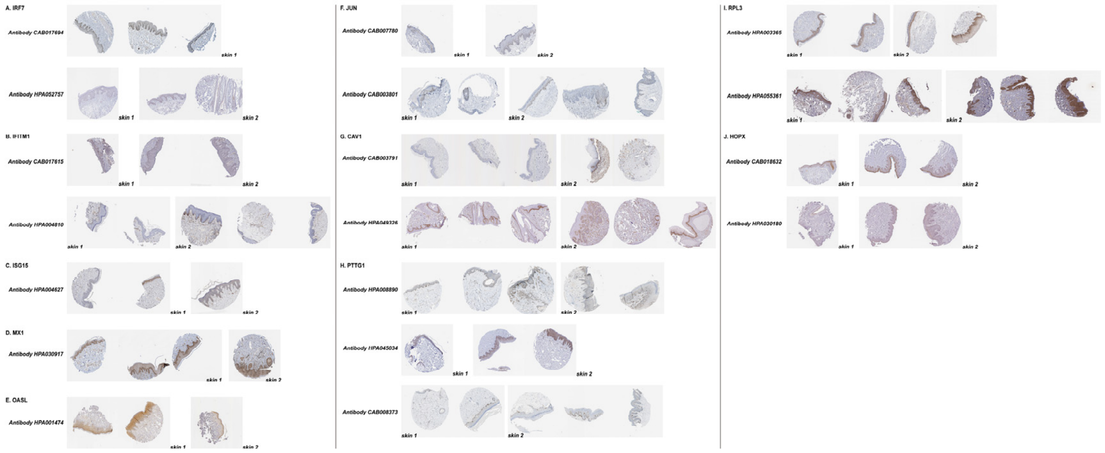

**Fig. S9**. Other valuable immunohistochemistry skin tissue microslides from HPA which can support cellular source evidence of UVBACGs. We suggest readers who are interested at these slides to check them in the website of HPA (https://www.proteinatlas.org/).

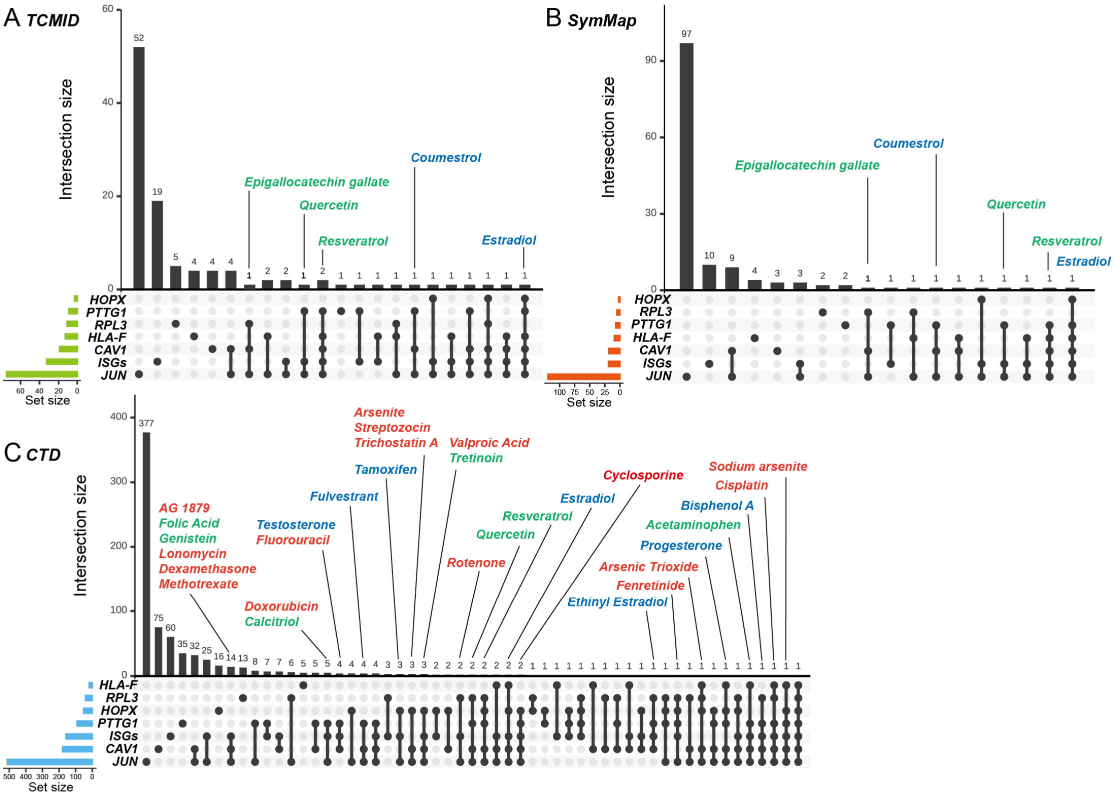


**Fig. S10.** Applying TCMID (Traditional Chinese Medicine Integrated Database, **A**), SymMap database (**B**), and CTD (Comparative Toxicogenomics Database, **C**) for identification of possible chemicals and herbal ingredients that might ameliorate disease activity by simultaneously regulating a variety of aberrant UVBACGs. Green, anti-inflammation chemicals; red, immunosuppressive or anti-tumor chemicals; blue, sex hormone-related chemicals. ISG: interferon-stimulated genes. The horizontal (green, orange, and blue) columns display the number of substances in the database that can regulate the corresponding UVBACG, while the vertical (black) columns show the number of substances in the database that can simultaneously regulate several UVBACGs, as indicated by the black dots in the diagram below.

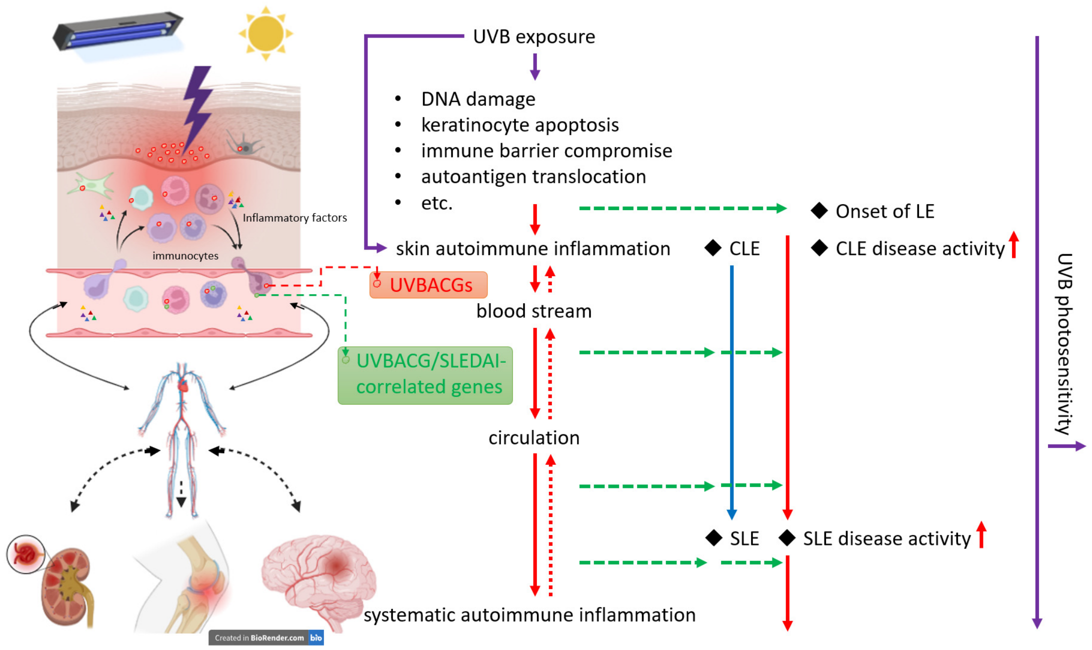


**Fig. S11. An illustrated summary of the key findings from this study.** UVB, as a crucial factor in LE diseases, can trigger the onset of LE, advance CLE to SLE, promote disease activity, and manifest SLE with CLE lesions. When enough area of the skin is exposed to a sufficient dose/time of UVB radiation, factors such as cell damage and autoantigen exposure, etc., drive the outbreak of skin inflammation, followed by autoimmune expansion, and systemic dissemination, with IFN-I playing a predominant role. Regarding photosensitivity of LE patients, to note, they have been carrying an existing autoimmune state (a pre-lesional state), where cellular UVBACGs, particularly ISGs, have already deviated from normal levels, lowering the threshold for the onset of skin and systemic inflammation. The more active the autoimmune cellular state in a patient, the lower the threshold for inflammation, indicating heightened photosensitivity. When the minimum dose of UVB exposure sufficient to trigger skin inflammation is measured, known as UVB-MED, it can be used to objectively quantify photosensitivity.
