## Supplementary figures and images for "Exploring the Correlation Between UVB Sensitivity and SLE Activity: Insights into UVB-Driven Pathogenesis in Lupus Erythematosus"

### Fig. S4

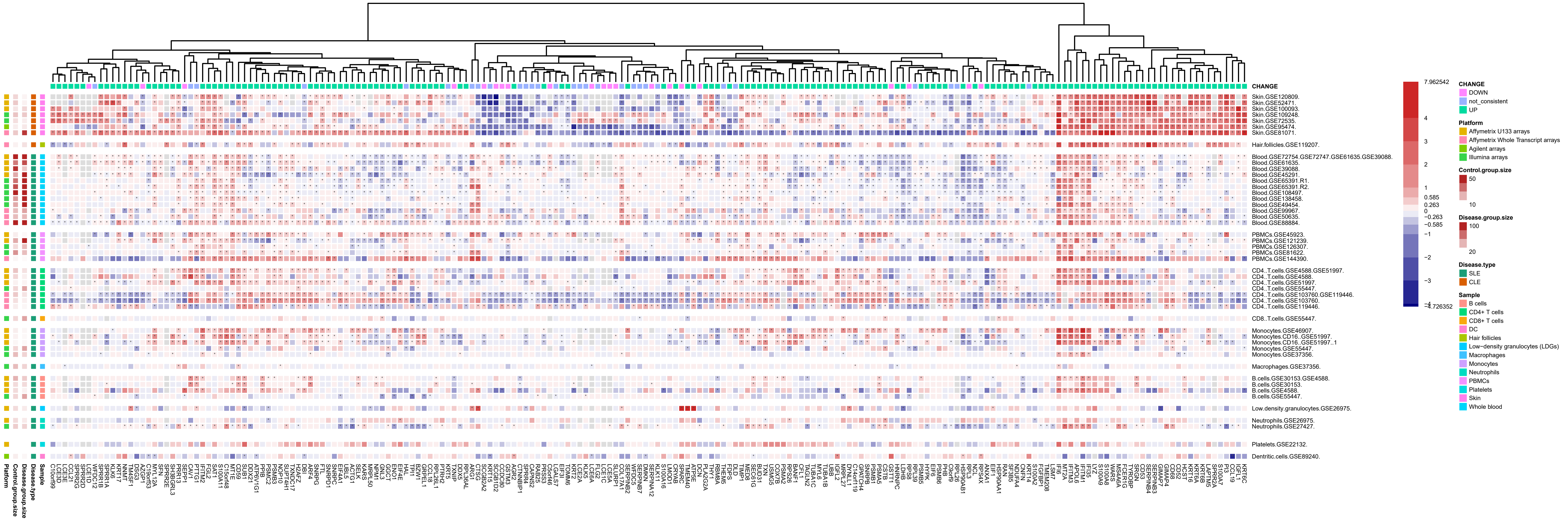

### Fig. S7

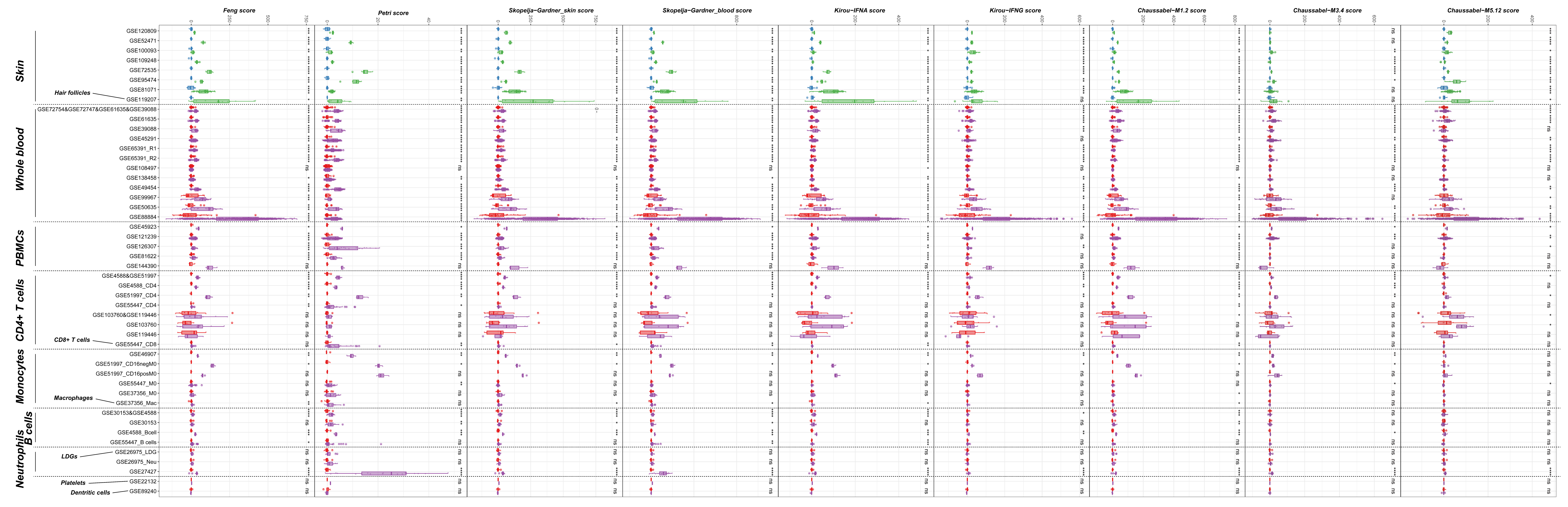
